# SFyNCS detects oncogenic fusions involving non-coding sequences in cancer

**DOI:** 10.1101/2023.04.03.535462

**Authors:** Xiaoming Zhong, Jingyun Luan, Anqi Yu, Anna Lee-Hassett, Yuxuan Miao, Lixing Yang

## Abstract

Fusion genes are well-known cancer drivers. However, very few known oncogenic fusions involve non-coding sequences. We develop SFyNCS with superior performance to detect fusions of both protein-coding genes and non-coding sequences from transcriptomic sequencing data. We validate fusions using somatic structural variations detected from the genomes. This allows us to comprehensively evaluate various fusion detection and filtering strategies and parameters. We detect 165,139 fusions in 9,565 tumor samples across 33 tumor types in the Cancer Genome Atlas cohort. Among them, 72% of the fusions involve non-coding sequences and many are recurrent. We discover two long non-coding RNAs recurrently fused with various partner genes in 32% of dedifferentiated liposarcomas and experimentally validated the oncogenic functions in mouse model.

## Background

Fusions between protein-coding genes caused by somatic SVs are well-known cancer drivers^1,2^, including *BCR*-*ABL1, EWS*-*FLI1, PML*-*RARA, TMPRSS2*-*ERG* and *FGFR3*-*TACC3*. It is estimated that 16% of cancers are driven by fusions^3^. Fusion proteins represent ideal drug targets since they do not exist in normal cells while tumor cell proliferation depends on them. One of the first targeted-therapy drugs in cancer, imatinib (Gleevec), is a small molecule inhibitor targeting the BCR-ABL1 fusion protein^4^. Many other inhibitors targeting different fusion proteins have since been approved for clinical use^5^. To date, more than 1,000 cancer-driving protein-coding fusions have been discovered^6^. However, only several oncogenic non-coding fusions have been reported, including *HERV-K*-*ETV1*^7^, *GAS5*-*BCL6*^8^, *USP9Y*-*TTTY15*^9^, *MALAT1*-*GLI1*^10^, *TTYH1*-*C19MC*^11^, *KDM4B*-*G039927* and *EPS15L1*-*lncOR7C2-1*^12^. A previous study on over 9,000 tumors from the Cancer Genome Atlas (TCGA) reported only 4% of fusions involving non-coding sequences^3^. This is because the algorithm used in that study, STAR-Fusion^3^, was designed to mainly detect protein-coding fusions, and therefore, the proportion of fusions involving non-coding sequences being 4% was certainly an underestimation. Fusions involving non-coding sequences are of clinical significance, as they can be used as biomarkers^13^ and studies are ongoing to target them therapeutically^14,15^. The discovery and characterization of the non-coding fusions may reveal new disease mechanisms and novel drug targets.

It is extremely challenging to differentiate true fusions from artifacts. Chimeric molecules in the sequencing library, sequencing errors, alignment errors and read-through fusions further complicate fusion detection. Most existing fusion callers depend on annotations of protein-coding genes and non-coding RNAs (ncRNAs), including DEEPEST^16^ and Arriba^17^. However, current ncRNA databases are still far from ideal because many ncRNAs are expressed at low levels and are highly tissue specific. The low expression also poses a major challenge to detect fusions involving non-coding sequences. Therefore, known oncogenic non-coding fusions remain rare. Another major roadblock is that a ground truth fusion set is not available, and most studies depend on in silico simulation, a small number of synthetic fusions, and validation on a small set of fusions to test the performances of the algorithms. Neither of aforementioned performance-testing strategies can be effectively used to comprehensively evaluate various fusion detection and filtering strategies and parameters. Here, we report a more sensitive computational algorithm “SFyNCS” to detect fusions involving non-coding sequences. We used somatic structural variations (SVs) detected from whole-genome sequencing data to validate fusions detected from RNAseq data. This allowed us to find the best performing fusion detection and filtering strategies. We then describe several recurrent and oncogenic fusions from 9,565 TCGA tumor samples. The oncogenic function of one of the recurrent fusions involving non-coding sequences was validated in mouse model.

## Results

### SFyNCS overview

Here, we developed <u>S</u>omatic <u>F</u>usions involving <u>N</u>on-<u>C</u>oding <u>S</u>equences (SFyNCS) to detect both protein-coding and non-coding fusions from RNAseq data (**Fig. 1a**). In this study, protein-coding fusions are defined as both fusion partners being protein-coding genes, whereas fusions involving non-coding sequences (FiNCS) have one or both fusion partners being non-coding sequences. We note that FiNCS may still encode proteins since the non-coding fusion partners may provide cryptic start or stop codons. SFyNCS searches for discordant read pairs and split reads, including those mapped to non-coding regions, to detect both protein-coding fusions and FiNCS (**Fig. 1b**). We use very loose cutoffs to detect raw fusions—one split read support required to define fusion breakpoints (**Methods**). Therefore, in the detection phase, SFyNCS is very sensitive, and a large number of raw fusions will be identified. Although many algorithms, such as STAR-Fusion^3^ and Arriba^12^, detect raw fusions similar to SFyNCS, the main advantage of SFyNCS lies in our search for the best performing filtering strategies (**Methods**). Since in silico simulations and synthetic fusions cannot fully mimic the artifacts and noise in real tumors, we sought to use fusions detected from real tumors to test fusion detection performances. Because ground truth fusions do not exist, to test performances, we took advantage of 338 tumor samples across 22 tumor types (**Supplementary Table S1**) with both RNAseq and whole-genome sequencing (WGS) data from the Cancer Genome Atlas (TCGA) cohort. Since tumor-specific fusions detected at the RNA level should be supported by somatic SVs detected at the DNA level, the 338 tumor samples allowed us to comprehensively evaluate different filtering strategies and cutoffs to determine the best performing filters. As it was not feasible to test all possible combinations of filtering strategies and cutoffs, we iteratively tested 49,248 combinations of cutoffs in three rounds (**Methods**) until no further improvement could be made (**Fig. 1c, 1d and Supplementary Table S2**). The final filters we chose to implement in SFyNCS with reasonable sensitivity and specificity were as follow: (1) at least one discordant read pair support; (2) at least one split read support; (3) at least three total read support (discordant read pair + split read); (4) the minimal distance between the discordant pairs and the split reads to be <=10 kb; (5) breakpoints for all intra-chromosomal fusions (deletion-like, duplication-like and inversion-like) not located in the same genes; (6) fusion breakpoint distance for deletion-like fusions to be >=500 kb; fusion breakpoint distance for duplication-like and inversion-like fusions to be >=20 kb; (7) standard deviation (SD) of fusion-supporting clusters within 100bp of breakpoints to be >=0.1; (8) canonical splicing motif present within 5bp of fusion breakpoints; (9) not found in any normal samples. The detailed description of the filters can be found in **Methods**. Using these filters, SFyNCS detected 12,923 fusions in the 338 samples (**Supplementary Table S3**) and 8,356 (64.7%) were supported by somatic SVs (**Fig. 2a**).

**Figure 1.**
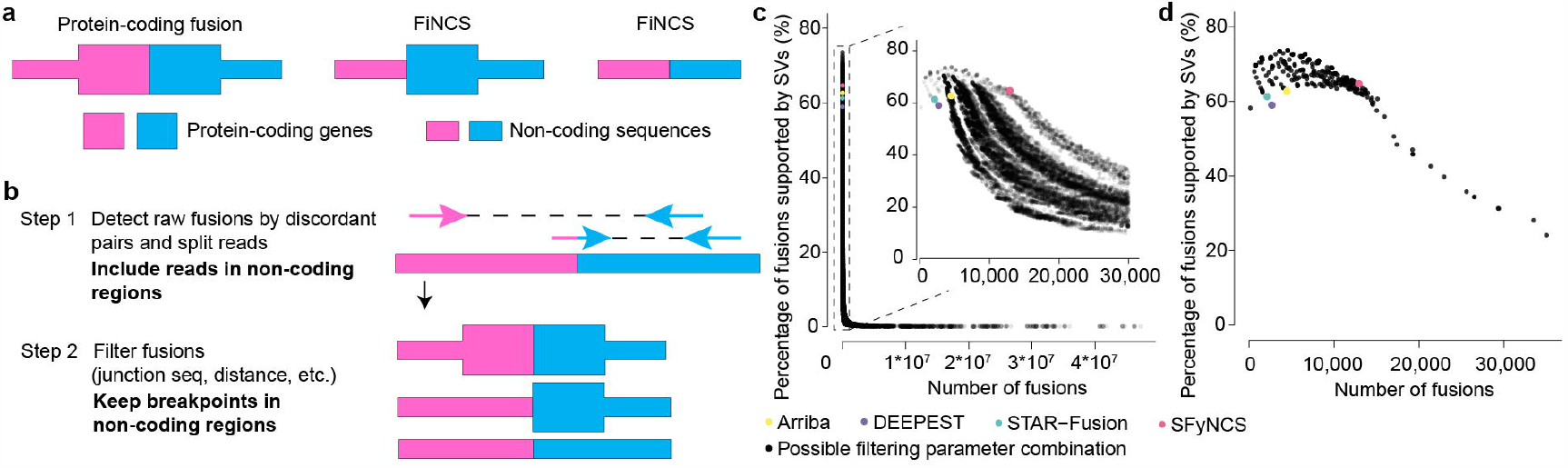
SFyNCS. **a**, Fusions of different types. Pink and blue shapes denote two fusion partners. Fusions can be in any combinations of protein-coding genes and non-coding sequences. **b**, Overview of SFyNCS. There are two main steps: detect raw fusions and filter fusions. **c**, A total of 49,248 combinations of filtering strategies and parameters are tested. Each dot represents one combination. The number of fusions is used to measure sensitivity and the percentage of fusions supported by somatic SVs is used to measure specificity. A portion of the plot is zoomed in in the upper right corner. **d**, Sensitivity and specificity of final filtering strategy implemented in SFyNCS compared to changing one parameter at a time. In both **c** and **d**, the sensitivity and specificity for Arriba, DEEPEST and STAR-Fusion are also shown.

**Figure 2.**
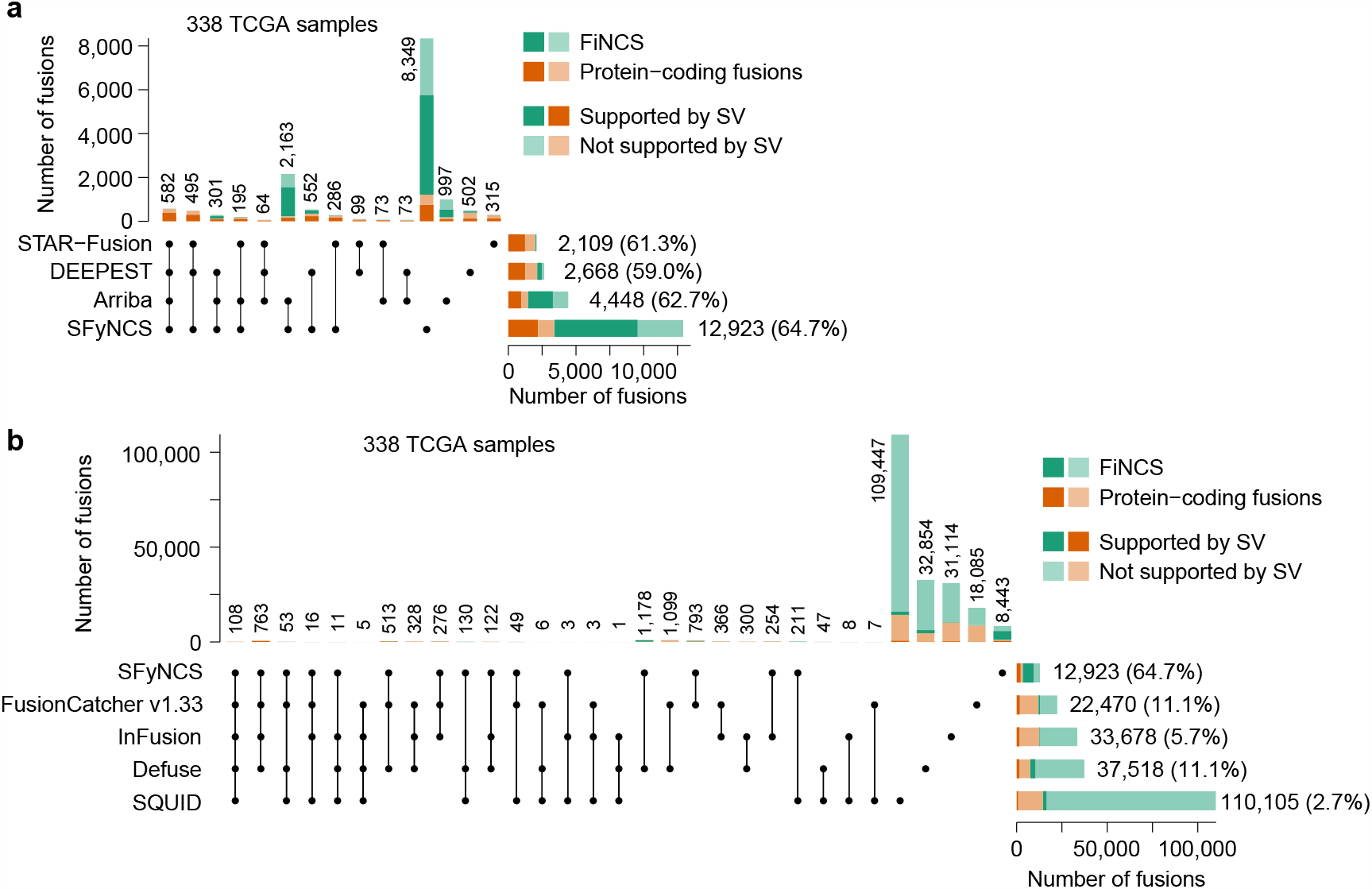
Benchmarking tools in TCGA samples. **a**, UpSet plot of four fusion-detection algorithms in 338 TCGA samples with both WGS and RNAseq data. The stacked bars on the bottom right are the total fusions detected by four tools respectively. The stacked bars on the top show the number of fusions identified by one or more tools. The black dots under the stacked bars indicate tools used. The numbers on the top and on the right side of the bars are numbers of fusions. The percentages in the parenthesis indicate percentages of fusions supported by somatic SVs. **b**, Comparison of SFyNCS with four fusion-detection algorithms, FusionCatcher v1.33, InFusion, Defuse, and SQUID, in the same 338 TCGA samples.

### Benchmarking SFyNCS

We compared SFyNCS with other algorithms in the same 338 samples from the previous section. Recently, STAR-Fusion^3^, DEEPEST^16^, and Arriba^12^ reported 2,109, 2,668 and 4,448 fusions in these samples, respectively (**Fig. 2a**). In contrast, SFyNCS detected 12,923 fusions which were 6.1, 4.8 and 2.9 folds of the ones detected by STAR-Fusion, DEEPEST, and Arriba, respectively. Therefore, the sensitivity of SFyNCS was far better than that of STAR-Fusion, DEEPEST, and Arriba. The fractions of fusions supported by somatic SVs were quite similar across the four algorithms, ranging from 59.0% to 64.7% (**Fig. 2a**). Fusions detected by SFyNCS had the highest SV support (64.7%). These suggested that the quality of fusions detected by these four algorithms were quite similar, and the specificity of SFyNCS was slightly better than that of STAR-Fusion, DEEPEST, and Arriba. Surprisingly, in the 12,923 SFyNCS-detected fusions, 9,520 (73.7%) were FiNCS. Among FiNCS, 64.7% were supported by SVs, which suggested that the quality of FiNCS detected by SFyNCS was as good as protein-coding fusions. STAR-Fusion and DEEPEST had limited ability in detecting FiNCS (**Fig. 2a**). Arriba detected 2,993 FiNCS and 2,145 of them were also detected by SFyNCS. SFyNCS detected 8,349 fusions that were missed by other algorithms and 63.3% of them were supported by SVs, which suggested that SFyNCS-specific fusions were of high quality. The vast majority (7,135) of these were FiNCS. In addition, SFyNCS detected 1,214 protein-coding fusions that were not detected by other algorithms. We then tested FusionCatcher^18^, InFusion^19^, Defuse^20^, and SQUID^21^ on the 338 tumors (**Supplementary Table S3**). These four algorithms detected many more fusions than SFyNCS, ranging from 22,470 to 110,105 (**Fig. 2b**). However, the fractions of fusions supported by SVs for these four algorithms ranged from 2.7% to 11.1% (**Fig. 2b**) indicating that the majority of these fusions were false calls. This suggested that the specificity of SFyNCS was far better than FusionCatcher, InFusion, Defuse, and SQUID.

We further tested SFyNCS on the breast cancer cell line MCF7 and compared to six algorithms that were previously tested^24^ on MCF7 (STAR-Fusion, MapSplice2^22^, InFusion, SOAPfuse^23^, FusionCatcher, and EasyFuse^24^). SFyNCS detected a total of 377 fusions including 262 (69.5%) FiNCS (**Fig. 3a** and **Supplementary Table S4**). In SFyNCS-detected fusions, 45.1% of the fusions were supported by SVs. STAR-Fusion, MapSplice2, InFusion, and SOAPfuse detected fewer fusions than SFyNCS (ranging from 70 to 256) and the fractions of fusions supported by SVs were lower than SFyNCS (ranging from 7.3% to 35.7%) (**Fig. 3a**). EasyFuse and FusionCatcher detected many more fusions (1,352 and 1,915 respectively). However, very few of them were supported by SVs (5.4% and 3.1% respectively) (**Fig. 3a**). In order to validate the fusions predicted by FusionCatcher, we extracted split reads provided by FusionCatcher and aligned them to the reference genome by BLAT. We found that only 16.5% of the fusions predicted by FusionCatcher were supported by the split reads, which was in sharp contrast to SFyNCS (80.6%) (**Supplementary Fig. S1a-S1e**). This suggested that the majority of fusions detected by FusionCatcher were likely false positives due to alignment errors. EasyFuse used 5 algorithms to detect fusions, including STAR-Fusion, MapSplice2, InFusion, SOAPfuse and FusionCatcher, and FusionCatcher was the only one detected a large number of fusions (**Fig. 3a**). Therefore, EasyFuse likely suffered from similar alignment errors. Among all these algorithms, only STAR-Fusion had comparable specificity to SFyNCS, but it detected five folds fewer fusions than SFyNCS. SFyNCS detected 275 fusions that were not detected by any other algorithms in MCF7 including 238 FiNCS. In the 275 SFyNCS-specific fusions, 49.1% were supported by SVs (**Fig. 3a**), which suggested that SFyNCS-specific fusions were of high quality. We randomly selected 20 FiNCS detected only by SFyNCS, performed PCR and Sanger sequencing validation, and were able to validate 12 (60%) of them (**Fig. 3b, Supplementary Fig. S2** and **Supplementary Table S5**). We further detected fusions in the MCF7 cell line using different RNAseq data produced by Cancer Cell Line Encyclopedia (CCLE) and Encyclopedia of DNA Elements (ENCODE) and found an additional 215 fusions (**Supplementary Fig. S1f** and **Supplementary Table S4**). We then randomly selected 10 FiNCS detected only in CCLE and ENCODE data and were able to validate 8 (80%) of them (**Fig. 3b, Supplementary Fig. S3** and **Supplementary Table S5**). Moreover, we validated 5 out of 6 (83%) randomly selected FiNCS in the colorectal cancer cell line HCT116 and the leukemia cell line K562 (**Fig. 3b, Supplementary Fig. S4, Supplementary Tables S5, S6** and **S7**).

**Figure 3.**
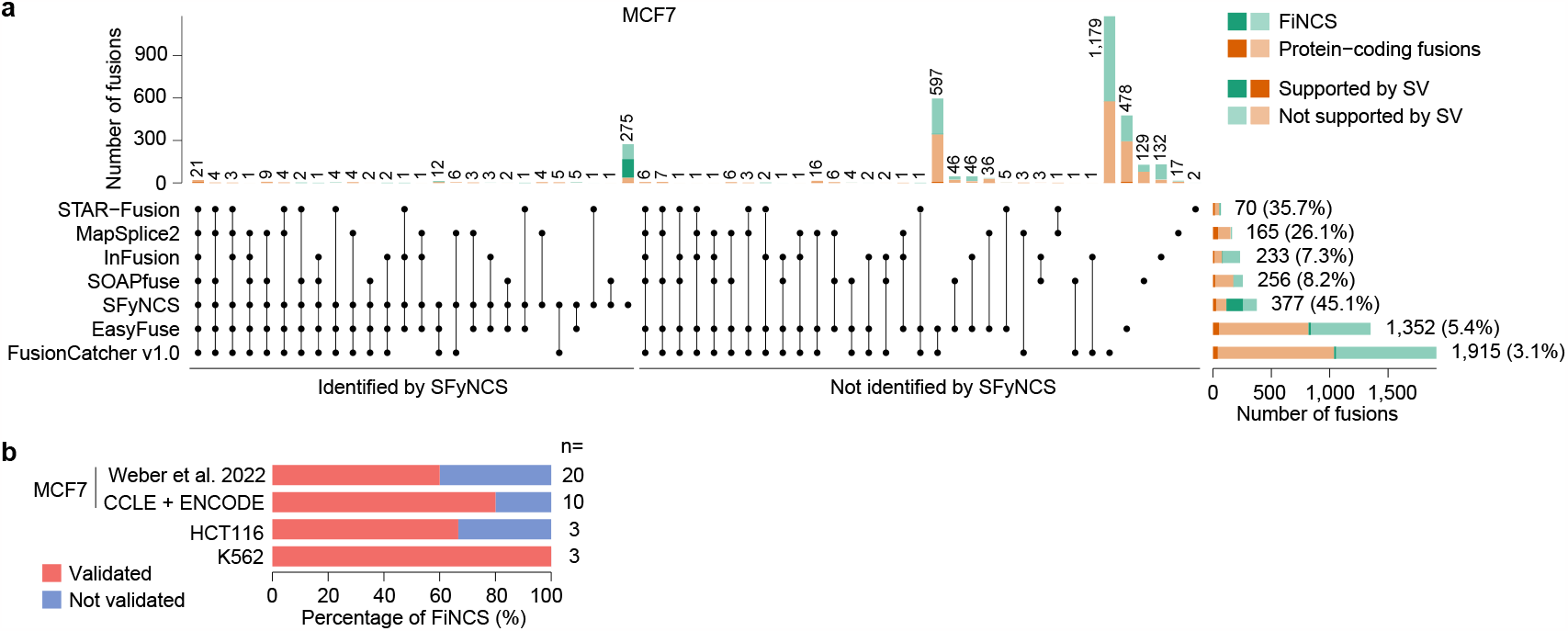
Benchmarking tools in MCF7 cell line. **a**, Comparison of SFyNCS with six fusion detection algorithms in MCF7 cell line: STAR-Fusion, MapSplice2, InFusion, SOAPfuse, EasyFuse, and FusionCatcher v1.0. Stacked bars on top are grouped into fusions identified by SFyNCS and not identified by SFyNCS. The stacked bars on the bottom right are the total fusions detected by seven tools respectively. The stacked bars on the top show the number of fusions identified by one or more tools. The black dots under the stacked bars indicate tools used. The numbers on the top and on the right side of the bars are numbers of fusions. The percentages in the parenthesis indicate percentages of fusions supported by somatic SVs. **b**, Percentages of FiNCS validated by PCR and Sanger sequencing in three cancer cell lines. The number of FiNCS tested are shown on the right side of bars.

Taken together, SFyNCS can detect many more fusions with better specificity than other existing algorithms, and the FiNCS detected by SFyNCS are highly accurate.

### Fusion landscape in TCGA cohort

We then used SFyNCS to analyze 9,565 TCGA tumor samples from 33 tumor types (**Supplementary Table S1**). A total of 165,139 fusions were detected (**Supplementary Tables S8**). Intriguingly, 119,191 (72.2%) of the fusions were FiNCS and were much more abundant than protein-coding fusions. Each tumor carried a median of 7 fusions ranging from 0 to 426 per tumor (**Supplementary Table S9**). Uterine Carcinosarcoma (UCS) and sarcoma (SARC) were the most abundant in fusions with medians of 32 and 29, respectively, whereas most kidney chromophobe cancers (KICH) and uveal melanomas (UVM) had less than 3 fusions (**Fig. 4a**). The abundance of fusions was consistent with somatic SV frequencies across tumor types^25^. STAR-Fusion, DEEPEST, and Arriba detected many fewer fusions in TCGA samples (25,664, 31,007 and 48,545, respectively)^3,12,16^. SFyNCS detected all known oncogenic fusions reported in these samples^3^ (**Fig. 4b**), such as *TMPRSS2*-*ERG, FGFR3*-*TACC3*, and *PML*-*RARA*. To better identify candidate driver FiNCS, we relied on recurrent fusion breakpoints at base-pair level since the annotation of non-coding genes remains incomplete. At the base-pair level, there were a total of 1,128 recurrent (occurring in at least 3 samples within the corresponding tumor type) fusion breakpoints involving non-coding sequences (**Fig. 4b, Supplementary Table S10**). Interestingly, except for prostate cancer (PRAD), the most recurrent fusion breakpoints involving non-coding sequences were often as frequent as protein-coding fusion breakpoints in many tumor types (**Fig. 4b**).

**Figure 4.**
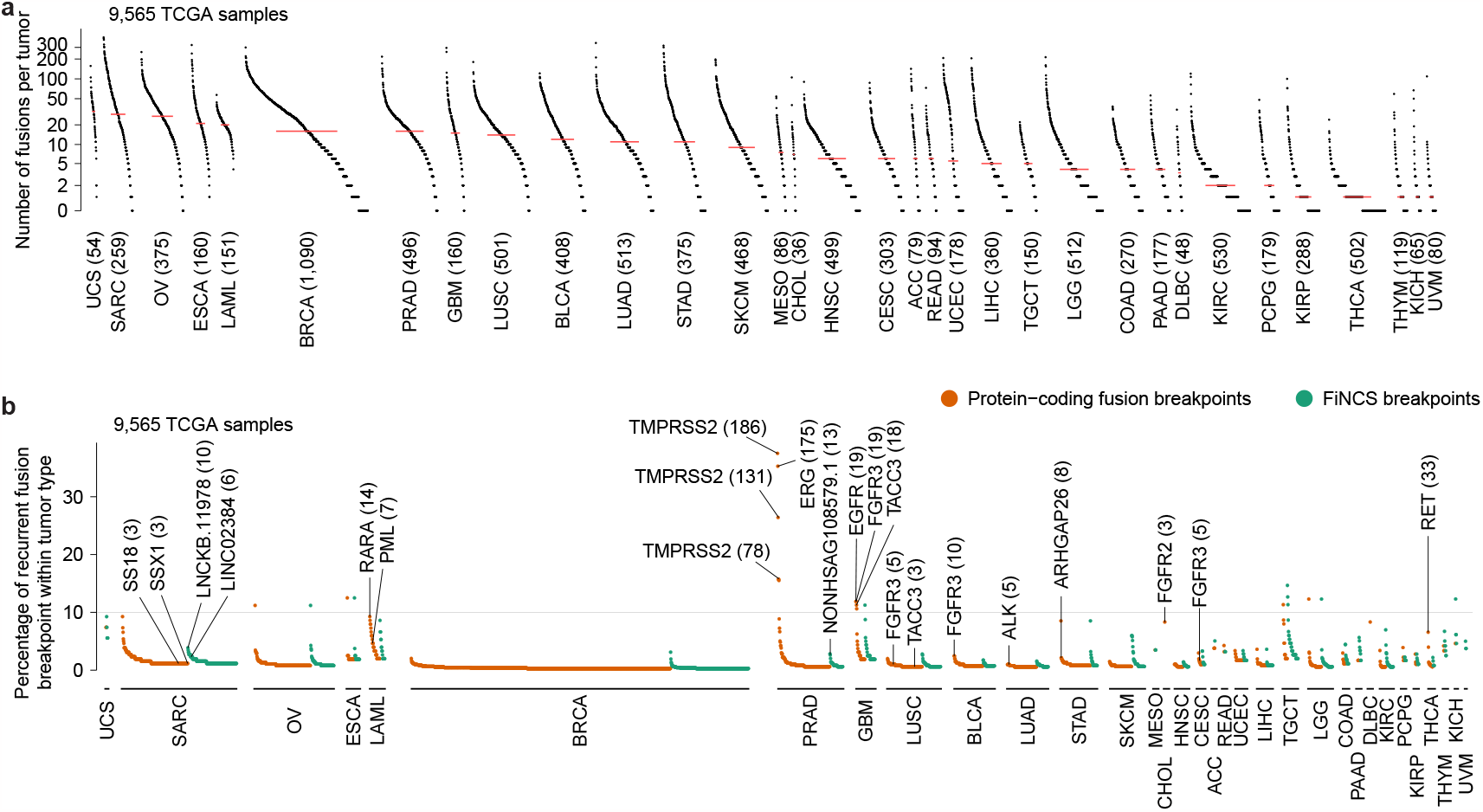
The landscape of fusion and recurrent fusion breakpoint in TCGA samples. **a**, The landscape of fusions in 9,565 TCGA samples. Each dot represents a tumor sample grouped by tumor type. Tumor types are sorted by median number of fusions per sample which is indicated by red lines. The numbers in the parenthesis are the numbers of tumor samples in the corresponding tumor types. **b**, Recurrent fusion breakpoints in 9,565 TCGA samples. Each orange or green dot represents a recurrent fusion breakpoint detected in at least three samples. The y axis indicates the percentage of samples carrying the fusion breakpoints in the corresponding tumor types. The numbers in parenthesis represent numbers of samples carrying the breakpoints. All breakpoints are at base-pair level. For example, *TMPRSS2*-*ERG* is the most recurrent fusion in adult solid tumors and can be detected in 183 out of 496 prostate cancers. Among them, 168 tumors have more than one *TMPRSS2*-*ERG* isoforms involving various exons of *TMPRSS2*. Therefore, 3 out of the top 4 recurrent fusion breakpoints in prostate cancer are in *TMPRSS2* gene and these breakpoints are observed in 186, 131 and 78 samples.

### Recurrent driver fusions involving non-coding sequences

In 496 prostate cancers, we identified 27 FiNCS in 13 samples (2.6%) involving a long non-coding RNA (lncRNA) on chromosome 17 *NONHSAG108579*.*1*. This lncRNA acted as the 5’ fusion partner (**Supplementary Table S11**). These FiNCS were mutually exclusive with the well-known ETS fusions (*P*=0.039, one-sided Fisher’s exact test, **Fig. 5a**). Two out of the 13 samples had WGS data, and in both samples, somatic translocations at the DNA level supported the FiNCS (**Fig. 5b** and **5c**). In sample TCGA-EJ-5518, there was a somatic translocation between chromosomes 8 and 17 (**Fig. 5b**). The translocation brought *NONHSAG108579*.*1* and *MYC* together and produced a chimeric transcript. Exons 2 and 3 of *MYC* were fused with *NONHSAG108579*.*1* and the chimeric transcript could produce an intact MYC protein (**Fig. 5b**). In another sample TCGA-CH-5771, there were two somatic translocations involving chromosomes 17 and 18 and resulting *NONHSAG108579*.*1* being fused to *ETV4* with an 8.9kb fragment from chromosome 18 inserted in-between (**Fig. 5c**). At the RNA level, the chromosome 18 fragment was entirely spliced out. On exon 9 of *ETV4*, there was an alternative start codon, and therefore, the *NONHSAG108579*.*1*-*ETV4* fusion transcript could produce a short ETV4 protein. The lncRNA *NONHSAG108579*.*1* was expressed at low levels in normal prostate tissues and fusion-negative prostate cancers, but highly expressed in most fusion-positive tumor samples (**Fig. 5b, 5c** and **Supplementary Fig. S5**). Most of the 3’ fusion partners were activated (**Fig. 5b, 5c**) and had expression patterns consistent with known driver fusions^26^, that is higher read coverage in exons included in the fusion transcripts than exons not part of the fusion transcripts. Furthermore, many of the 3’ fusion partners were well-known oncogenes including *MYC, ETV4, ETV1* and *BRAF* (**Supplementary Table S11**). Therefore, the *NONHSAG108579*.*1* fusions in prostate cancers were highly likely to be oncogenic.

**Figure 5.**
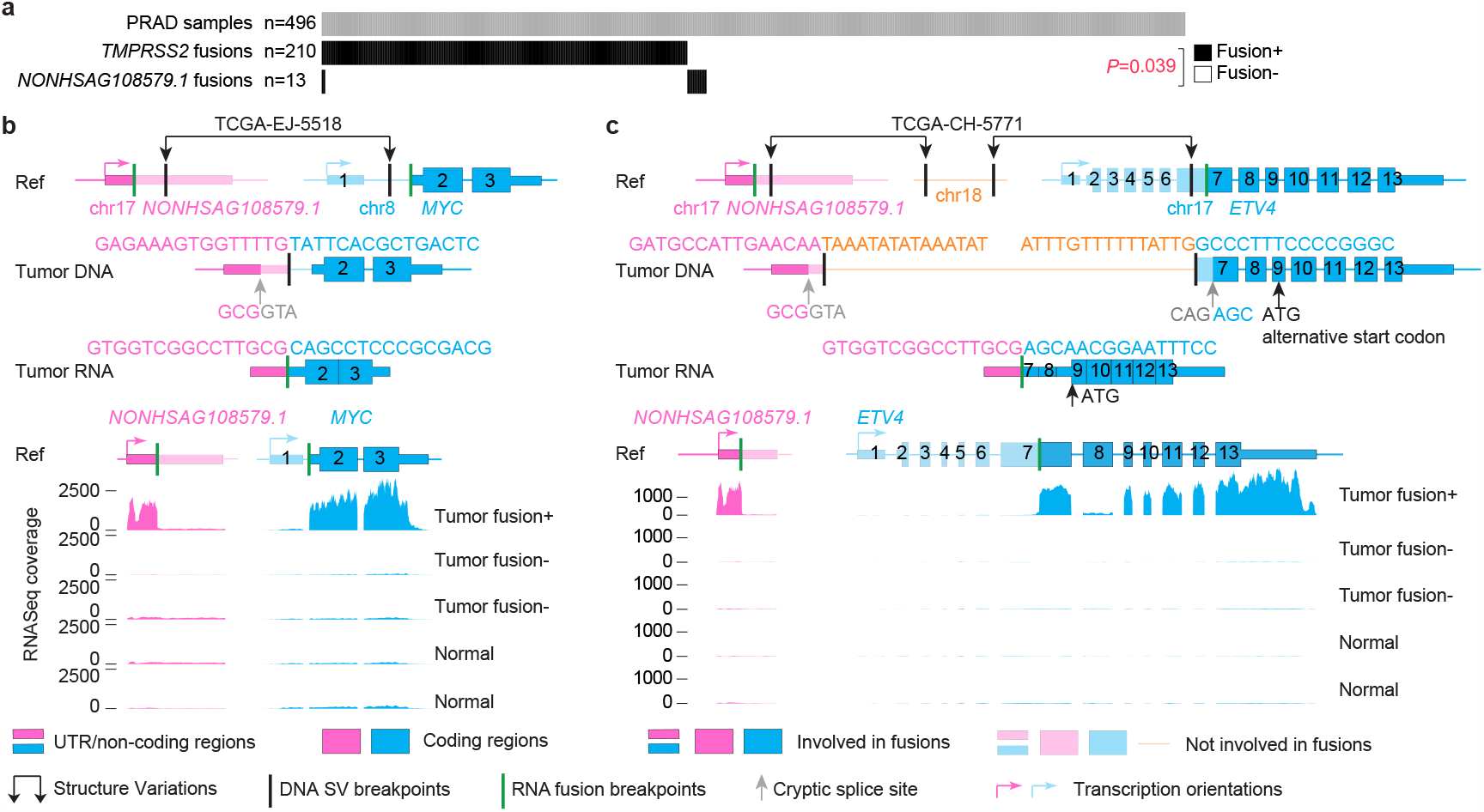
Recurrent FiNCS in prostate cancer. **a**, Oncoprint plot of 496 prostate cancers showing fusions involving *TMPRSS2* and *NONHSAG108579*.*1*. **b** and **c**, Structures of two *NONHSAG108579*.*1* fusions and their expression. The top three rows are gene and fusion structure cartoons of the reference genome, tumor DNA, and tumor RNA. Pink and blue boxes denote two fusion partners. The *NONHSAG108579*.*1*-*ETV4* fusion in sample TCGA-CH-5771 is produced by two different translocations. The orange fragment from chromosome 18 is entirely spliced out from the fusion transcript. Five tracks of RNAseq coverage are shown for five samples at the bottom and the reference gene structures are given above the five tracks. Exons and introns are re-scaled to better illustrate fusion structures. In **b**, the tumor samples without fusions (fusion-) are TCGA-HI-7169-01A-11R-2118-07 and TCGA-EJ-A7NJ-01A-22R-A352-07, the normal samples are TCGA-EJ-7327-11A-01R-2118-07 and TCGA-HC-7742-11A-01R-2118-07. In **c**, the fusion-samples are TCGA-G9-6365-01A-11R-1789-07 and TCGA-HI-7169-01A-11R-2118-07, the normal samples are TCGA-EJ-7123-11A-01R-1965-07 and TCGA-EJ-7125-11A-01R-1965-07.

In addition, recurrent FiNCS involving two lncRNAs (*LINC02384* and *LNCKB*.*11978*) were detected in 259 sarcomas (**Supplementary Table S12**). All of these FiNCS were detected in dedifferentiated liposarcomas (DDLPS), not other subtypes, and they were mutually exclusive with each other (**Fig. 6a**). *LINC02384* and *LNCKB*.*11978* fusions occurred in 6 (12%) and 10 (20%) DDLPS tumors, respectively, and both lncRNAs were the 3’ fusion partners. The 5’ fusion partners were either protein-coding genes, lncRNAs or pseudogenes (**Supplementary Table S12**). Among the 16 fusion-positive tumors, 6 had WGS data, and somatic SVs at the DNA level supported the FiNCS in all 6 samples (**Fig. 6b, 6c, Supplementary Fig. S6** and **S7**). In sample TCGA-DX-A1L3, a somatic tandem duplication was present in protein-coding gene *ZDHHC17* and upstream of *LNCKB*.*11978* (**Fig. 6b**). Exon 1 of *LNCKB*.*11978* was skipped and a chimeric transcript of exon 1 of *ZDHHC17* and exon 2 of *LNCKB*.*11978* was produced. The transcript could be translated into *LNCKB*.*11978* and produced a chimeric protein (**Fig. 6b**). In sample TCGA-DX-A3LY, there was a somatic translocation between chromosomes 5 and 12 (**Fig. 6c**). Similarly, a transcript of exon 1 of *SH3RF2* and exon 2 of *LINC02384* was produced and could be translated into a chimeric protein (**Fig. 6c**). In most of these FiNCS involving *LNCKB*.*11978* and *LINC02384*, the 3’ lncRNAs were activated (**Fig. 6b, 6c, Supplementary Fig. S6** and **S7**). The high recurrence and expression patterns indicated that these FiNCS were potential cancer drivers. To test the oncogenic functions experimentally, we synthesized the *ZDHHC17*-*LNCKB*.*11978* fusion, transduced it into A549 cells (**Fig. 6d**), and injected the cells into immune deficient mice subcutaneously. Although the cancer cells don’t grow differently in culture, tumors carrying the fusion grew significantly faster than controls (**Fig. 6e** and **6f**) upon grafting on mice, suggesting that the *ZDHHC17*-*LNCKB*.*11978* fusion does indeed have oncogenic activity.

**Figure 6.**
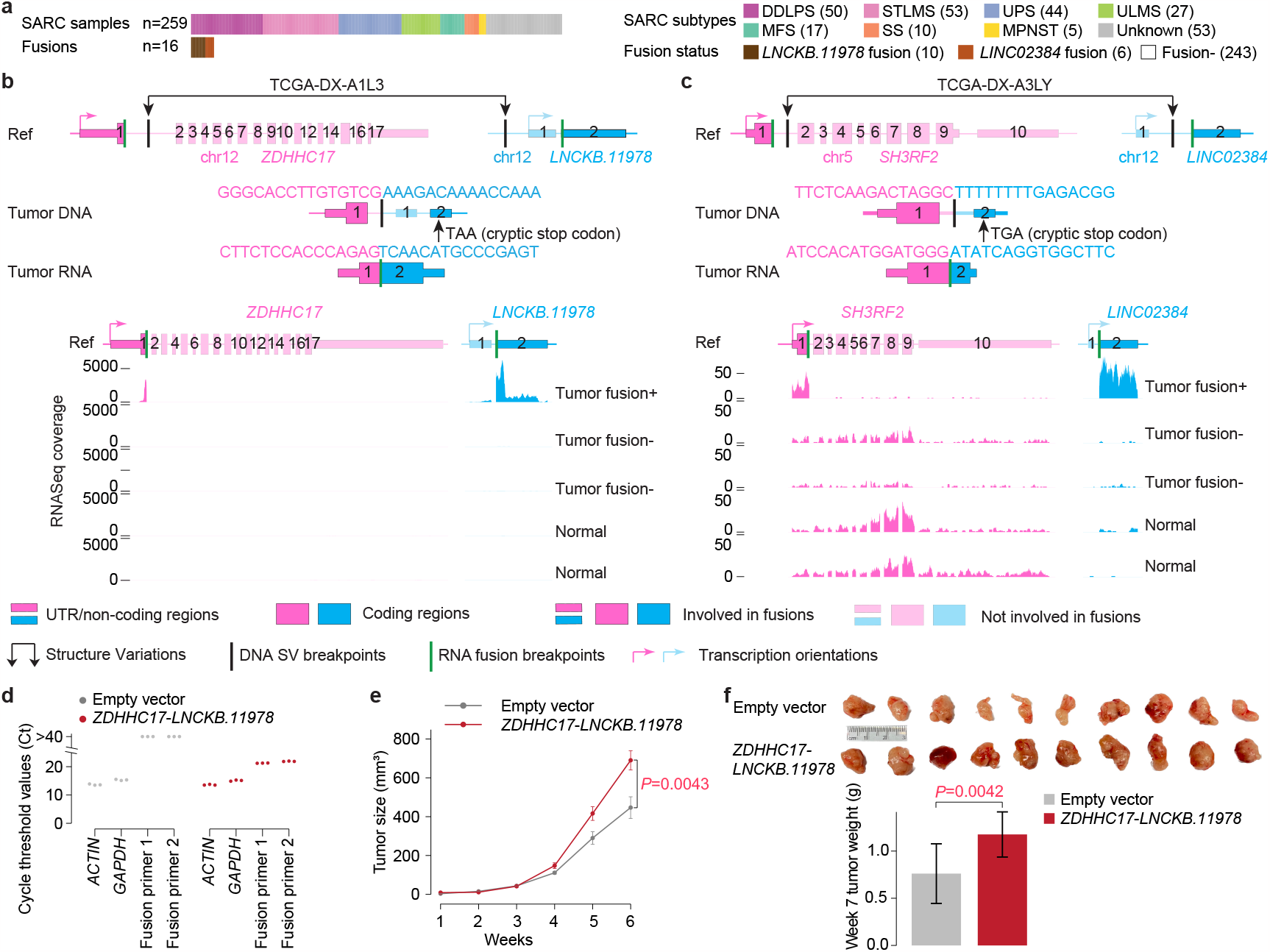
Recurrent FiNCS in sarcoma. **a**, Oncoprint plot of 259 sarcomas showing FiNCS involving *LNCKB*.*11978* and *LINC02384*. DDLPS: dedifferentiated liposarcoma, STLMS: Soft Tissue Leiomyosarcoma, UPS: Undifferentiated Pleomorphic Sarcoma, ULMS: Gynecologic Leiomyosarcoma, MFS: Myxofibrosarcoma, SS: Synovial Sarcoma, MPNST: Malignant Peripheral Nerve Sheath Tumor. **b** and **c**, Structures of a *LNCKB*.*11978* fusion and a *LINC02384* fusion in DDLPS and their expression. The top three rows are gene and fusion structure cartoons of the reference genome, tumor DNA, and tumor RNA. Pink and blue boxes denote two fusion partners. The tumor samples without fusions (fusion-) are TCGA-IE-A4EI-01A-11R-A24X-07 and TCGA-IW-A3M4-01A-11R-A21T-07, the normal samples are SRX636240 and SRX640265 respectively. **d**, Quantitative PCR showing the presence of *ZDHHC17*-*LNCKB*.*11978* fusion transcript in A549 cells. **e**, Tumor growth curves after subcutaneous injection from week 1 to week 6. Error bars are standard deviations. *P* value is calculated by two-sided Student’s t-test. **f**, Pictures of 10 tumors and tumor weights at week 7 after subcutaneous injection. Error bars are standard deviations. *P* value is calculated by two-sided Student’s t-test. t-test. **f**, Pictures of 10 tumors and tumor weights at week 7 after subcutaneous injection. Error bars are standard deviations. P value is calculated by two-sided Student’s t-test.

Taken together, our results demonstrate that SFyNCS is able to detect oncogenic fusions involving non-coding sequences.

## Discussion

Here, we describe our fusion detection algorithm SFyNCS which can detect fusions of both protein-coding genes and non-coding sequences in transcriptome sequencing data. SFyNCS is designed for Illumina short-read sequencing data and will suffer from the limitations of short-read sequencing technology, such as the lack of ability to resolve repetitive regions since human genome is highly repetitive. Fusion breakpoints in transposable elements, segmental duplications, satellite repeats, simple repeats and other types of repeats are unlikely to be reliably detected. This constraint is not specific to SFyNCS. All short-read based fusion detection algorithms suffer from this limitation.

Another obstacle is the availability of normal samples to filter out germline events and systematic artifacts. Several tumor types do not have RNAseq data from matched normal samples, such as acute myeloid leukemia (LAML), lower grade glioma (LGG), ovarian cancer (OV,) testicular germ cell tumors (TCGT), uterine carcinosarcoma (USC), while some tumor types have very few matched normal samples, such as esophageal cancer (ESCA), glioblastoma (GBM), skin cutaneous melanoma (SKCM), thymoma (THYM). Therefore, many of the highly recurrent fusions detected from these tumor types are likely not cancer drivers.

Although SFyNCS displayed superior performances in our benchmarking tests compared to existing tools, a small fraction of true fusions were still missed by SFyNCS. Each filter we implemented may remove some true fusions, such as true fusion junctions may not always be canonical splice sites^26^. For other types of somatic variants including single nucleotide variants (SNVs), copy number variations (CNVs) and SVs, multiple tools are often integrated together for variant calling^27^. Therefore, we recommend users to apply multiple tools to perform comprehensive fusion detection.

## Conclusion

We report our tool SFyNCS to detect fusions involving non-coding sequences. With rigorous benchmarking using tumor samples and cancer cell lines, we show that SFyNCS is more sensitive in fusion detection than existing tools and the quality of fusions detected by SFyNCS is better than existing tools. About three quarters of the fusions in tumor samples have non-coding fusion partners. Some recurrent fusions involving non-coding sequences can promote tumorigenesis.

## Methods

### SFyNCS Workflow

#### Identifying raw fusions

RNAseq reads were aligned by STAR^28^ to the reference genome for detection of discordant read pairs and split reads. Discordant pairs defined by STAR were paired-end reads aligned to different chromosomes or to the same chromosome but in incompatible orientations, or in compatible orientations but with distances greater than 100 kb. Some reads could not be aligned consecutively in the genome but had to be split into two parts. If the two parts were aligned to two different chromosomes or to the same chromosome but in incompatible orientations, or in compatible orientations but with distances greater than 100 kb, these reads were considered split reads which potentially spanned the fusion breakpoints. Discordant pairs and split reads aligned to multiple locations were discarded and duplicated reads (read pairs with identical mapping) were removed. Discordant pairs and split reads were merged into clusters if they were aligned to the same chromosomes, with the same orientations and within 1 Mb to each other. Raw fusions were then called from these clusters. Precise fusion breakpoints were determined by split reads. Split reads with same orientations and within 5bp were considered to support the same fusion. Each candidate fusion must be supported by at least one split read. In the initial detection phase, discordant read pair support was not required. Different numbers of read support (discordant read pair and split read) were tested in a later section. Note that one discordant pair may support more than one fusion (different isoforms) depending on how the transcripts were spliced (**Supplementary Fig. S8**). Gene annotation was not used in raw fusion detection, so that fusion breakpoints in both protein-coding genes and non-coding regions of the genome could be detected. The process described above was very sensitive, and hence, a large number of raw fusions would be detected in each sample.

#### Testing filtering strategies

To detect high quality tumor-specific fusions, we comprehensively tested the performances of the fusion calling and filtering strategies as well as various cutoffs in three rounds. In the first round, we intended to find what filters were useful and tested the following: (1) Number of total read support (discordant pair and split read combined, cutoffs tested: >=2 and >=3); (2) Number of split read support (cutoffs tested: >=1 and >=2); (3) Number of discordant pair support (cutoffs tested: 0 and >=1); (4) The minimal distance between the discordant pairs and the split reads supporting the same fusion (cutoffs tested: <=5 kb, <=10 kb and NA [filter not applied]); (5) Whether filter deletion-like fusions that were within the same gene annotated by GENCODE or not; (6) Whether filter duplication-like and inversion-like fusions that were within the same gene annotated by GENCODE or not; (7) Fusion breakpoint distance for deletion-like fusions (produced by somatic deletions at the DNA level, cutoffs tested: >=200 kb and >=500 kb and NA); (8) Fusion breakpoint distance for duplication-like and inversion-like fusions (produced by somatic duplications and inversions at the DNA level, cutoffs tested: >=10 kb, >=20 kb and NA); (9) Breakpoint flanking sequence identity by aligning 20bp sequences (10bp from both sides) of two breakpoints with Needleman–Wunsch algorithm (cutoffs tested: <=0.5 and NA); (10) Size of breakpoint flanking region for filters (11) and (12) (cutoffs tested: 100bp); (11) Standard deviation (SD) of fusion-supporting read clusters in fusion breakpoint flanking region (described in detail in the next paragraph, cutoffs tested: >=0.05, >=0.1 and NA); (12) Number of fusion-supporting clusters in fusion breakpoint flanking region (cutoffs tested: <=5 and NA); (13) Filter by canonical splicing motifs (GT in the donor site, AAG/CAG/TAG in the acceptor site) within 5bp of fusion breakpoints; (14) Confirming discordant pairs and split reads alignment by TopHat2 (distance between TopHat2 and STAR alignments of split reads <=5bp); 15) Confirming split reads alignment by BLAT; and (16) Filter by fusion breakpoints detected in normal samples (more details below).

In the second round, to optimize the filtering parameters, we further tested more cutoffs based on the results from the first round by changing one or a few parameters at a time: (1) Number of total read support (cutoffs tested: >=3, >=4 and >=5); (2) Number of split read support (cutoffs tested: >=2, >=3, >=4 and >=5); (3) The minimal distance between the discordant pairs and the split reads supporting the same fusion (cutoffs tested: <=100bp, <=200bp, <=500bp, <=1kb, <=5kb, <=20kb, <=50kb, <=100kb, <=200kb, <=300kb, <=500kb, <=1Mb and NA); (4) Fusion breakpoint distance for deletion-like fusions (cutoffs tested: >=100kb, >=200kb, >=300kb, >=500kb, >=1Mb and NA); (5) Fusion breakpoint distance for duplication-like and inversion-like fusions (cutoffs tested: >=10kb, >=30kb, >=50kb, >=100kb, >=200kb, >=300kb, >=500kb, >=1Mb and NA); (6) Breakpoint flanking sequence identity (cutoffs tested: <=0.3, <=0.5 and <=0.8); (7) Different size of breakpoint flanking region in (8) and (9) (cutoffs tested: 100bp, 500bp, 1kb, 5kbp and 10kb); (8) SD of fusion-supporting read clusters in fusion breakpoint flanking region (cutoffs tested: >=0.05, >=0.15, >=0.2, >=0.25, >=0.3 and NA); (9) Number of fusion-supporting clusters in fusion breakpoint flanking region (cutoffs tested: <=5, <=10, <=15, <=20, <=25, <=30 and NA); Note that, in the second round, not all possible parameter combinations were tested. A selected subset based on the best performing combination from the first round were tested to find better performing parameters.

In the third round, we either removed one filter, added one filter, or changed the cutoff for one filter based on the best performing filter combination determined in the second round to confirm that no further improvement could be made (**Supplementary Table S2**).

For each candidate fusion breakpoint, there could be more than one read cluster supporting different fusions in its flanking region. Too many such clusters suggested that the alignments of this region were unreliable. The number of fusion-supporting clusters was tested. Standard deviations (SDs) of the proportions of fusion-supporting reads in these clusters (equation below) was tested.

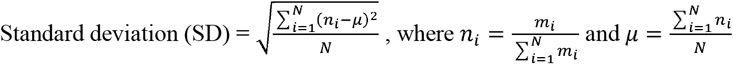

N is the number of clusters, m_i_ is the number of reads in cluster i, n_i_ is the proportion of reads in cluster i.

Normal samples from TCGA (**Supplementary Table S13**) were used to remove germline events and other systematic artifacts. A panel of 140 normal samples was first constructed by randomly selecting 10 normal samples from each tumor type that had more than 10 matched normal samples. Fusions detected in each tumor sample were filtered by this normal panel as well as all the matched normal samples of the corresponding tumor type when available. Note that some tumor types, such as lower-grade glioma and ovarian cancer, did not have matched normal samples. These tumor samples were only filtered by the 140-sample normal panel. Fusions detected in tumor samples were discarded if there were at least two fusion supporting reads (either discordant read pairs or split reads) within 10 kb for both breakpoints in any normal samples.

Note that if the fusion breakpoints were located close to the end of the transcripts, discordant read pairs may not exist. Therefore, we tested the fusion detection performance if not requiring discordant read pair support. Since fusion breakpoints were determined by split reads, we did not test fusion detection without split read support.

### Benchmarking fusion detection tools

Fusions in 338 TCGA samples were identified by Defuse (v0.8.1), FusionCatcher (v1.33), InFusion (v0.8.1-dev), and SQUID (v1.5) with default parameters. Note SQUID failed to analyze TCGA-DX-A2IZ-01A-11R-A21T-07. Fusions detected by multiple tools needed to have identical breakpoint locations and orientations. Fusions were considered supported by somatic SVs if SV breakpoints could be found within 100 kb of fusion breakpoints and the DNA fragments produced by the SVs could be spliced into the corresponding fusion RNA. Fusions in MCF7^24^ were identified by FusionCatcher (v1.33) with default parameters. Fusion-supporting split reads identified by both FusionCatcher (v1.33) and SFyNCS were aligned to the reference genome by BLAT to validate split-read alignment. If there were two segments of a split read aligned uniquely within 5bp of the predicted fusion breakpoints, the split read was considered validated by BLAT. Split reads not validated by BLAT mainly belonged to the following three categories: i) align entirely (more than 85bp of 101bp-long reads) to one location of the genome (**Supplementary Fig. S1c**), ii) not support (aligned within 5bp of the predicted breakpoints) one or more fusion breakpoints (**Supplementary Fig. S1d**), or iii) align to multiple locations (**Supplementary Fig. S1e**). If a fusion did not have any split read validated by BLAT, the fusion was considered not validated.

### Cell lines

HEK293T cells were obtained from Dr. Alexander Muir (University of Chicago). MCF7 cells were obtained from Dr. Lev Becker (University of Chicago). HCT116 and K562 cells were obtained from Dr. Chuan He (University of Chicago). A549 cells were purchased from ATCC (American Type Culture Collection, USA). All cell lines were cultured at 37°C/5% CO2. HEK293T cells were cultured in Dulbecco’s Modified Eagle Medium (DMEM) (Gibco, 21041025) supplemented with 10% FBS, 1% penicillin/streptomycin and 2 mM L-glutamine. MCF7 cells were cultured in Eagle’s Minimum Essential Medium (Corning, 10-010-CV) with 10% fetal bovine serum (FBS) (Gibco, A4766). HCT116 cells were cultured in McCoy’s 5A Medium Modified (Gibco, 16600-082) with 10% FBS. K562 cells were cultured in Iscove’s Modified Dulbecco’s Medium (Gibco, 12440-053) with 10% FBS. A549 cells were cultured in F-12K Medium (ATCC, 30-2004) with 10% FBS and 1% penicillin/streptomycin. All cell lines have been regularly monitored and tested negative for mycoplasma using the mycoplasma detection kit (Lonza, LT07-218).

### RT-PCR and Sanger sequencing validation

Twenty fusions were randomly selected for validation among the 238 FiNCS in MCF7 RNA-seq data^24^ detected by SFyNCS but not detected by FusionCatcher (v1.0), InFusion (v0.8), MapSplic2 (v2.2.1), SOAPfuse (v1.2.7), STAR-Fusion (v1.5.0), or EasyFuse (v1.3.0). Ten FiNCS detected in MCF7 RNA-seq data produced by CCLE and ENCODE but not detected in the RNA-seq data produced by the previous study^24^ were randomly selected. Six FiNCS were randomly selected from HCT116 and K562 cell lines. Primers (**Supplementary Table S5**) were designed by Primer3 and synthesized by Integrated DNA Technologies. MCF7, HCT116 and K562 cells were plated in 6-well plates and allowed to reach 80% confluence prior to RNA extraction. After cells being lysed in 300µl/well TRYzolTM (Invitrogen, 15596026), RNA samples were prepared following the manual of Direct-zol RNA Miniprep kit (RPI, ZR2052). Reverse transcription was performed using Applied Biosystems High-Capacity cDNA Reverse Transcription Kit (43-688-14) following manufacturer’s instructions. PCR was conducted on SimpliAmpTM Thermo Cycler (Applied Biosystems, A24811), with HotStarTaq Plus Master Mix (QIAGEN, 1039620) following the manufacturer’s instructions. PCR products were extracted from 2% agarose gel with MinElute Gel Extraction kit (QIAGEN, 28604) and purified with MinElute PCR purification kit (QIAGEN, 28004). Then the DNA samples were sent to the DNA Sequencing & Genotyping Facility of the University of Chicago Comprehensive Cancer Center for Sanger sequencing.

### Synthesis of *ZDHHC17-LNCKB*.*11978*.*4*

The 1,870 bp *ZDHHC17-LNCKB*.*11978*.*4* fusion cDNA was synthesized by GenScript (New Jersey, USA) and subcloned into the lentiviral pCDH-CMV-MCS-EF1-Puro plasmid (SBI, CD510B-1). The cDNA sequence in the plasmid was verified by Sanger sequencing at University of Chicago Medicine Comprehensive Cancer Center core facility. The *ZDHHC17-LNCKB*.*11978*.*4* fusion cDNA sequence is TTGTATCCATGTTTTTCCGGGCGTCCCCCGGAGGGACAGGTTGCGGGTGACCTTTTCAAGTGTGGAGGAAAGGGAAGCTGCTTTTGTCTTCAGGAATGATGCAGGTCTCGACTCAAGCCTGACGGGCCCAAACCTCCCTGGAGCTGGCTGACGACTCTGCCCGAGTTCCTGAAGAGGGGTCCCGGGGGTCCCGGAGCGGAAGTGGGAGCGCGTGGGCGTGGGCTCCTCGGCTGCCTGGGGCTCCAGACTTGTGCTGCGTGCGGCTCCGGAGCTCTGTTCTCGCTCCTGAGCAGCTGCTAGGTTTCCCAAGCGACTGTCTCAACCGCCCGGCCGCCTCCCCCGGGCAGCCAGAGCTTCACATCTACCTCCAGCCGGGACCCGCCCCCGAGCCGCGGGGCCCACGCCCAGAGCCCTCCGCCGTCCCCAGCGCAGTGCAGCAGAGCGCGATCCAGTCTGGGGCCGGGCCGCGCTTCCGCGCACGCGCGGAGAAACCCGCGCCCTCCGAGGGG GGAGGGGACAGAGGGGGCGTCACGGGGGCAGGAGAAGAAGGAGGAGGAGGCCCGCGTCGCCTCCGGCGGGGCTCGCGCTCGCCCCGCGCTCGCCCTCCGCCTCGCCCGAGCCCCGGGAGGGTGAAACGCTTTCTCCCAGCATGCAGCGGGAGGAGGGATTTAACACCAAGATGGCGGACGGCCCGGATGAGTACGATACCGAAGCGGGCTGTGTGCCCCTTCTCCACCCAGAGTCAACATGCCCGAGTGCTGTGAACGTTATGAGAGGGCCTTGTTGGGAACACGTGCTCCTGGGAATCAGCCCTTCCCTCTGTCCTGTTCCCACTCCTCCCCGACGATGCTCCTGCTCAGAACCCACTCCTCACCTCAGTGAAGCAACGCAGCGGGCACCCTGTGGACAAAGCTGGATATTGGCTCTGAATAAAAGCGAATCATGGGGAAAATCAGTGTCTCAGTAAAATGGGGTTTTCTTAGTAGAGACCAGACTGTGAAGGACCTTGCTTCATTCCATCTTTGAGGAGGATGATGATTCAGGGACATTGGCCCAAGATCAAAGTGGTATTTTTAGGTTGTATTTACTTAGCTATTTGCCGTCTACCTCCTTATTTCCAGGTAGCAACTTCCTTCTTATATCTGAGATGTTTAAGAGATGATGAAACCAGCTTGCACACACTTCTCAAAGTGTGTTTGTTCGCATCCATTATTTCACTGGGGACCGGCTATTATCCTCTCCATTTTCTTTATAAGGATATTGAAAGAGAGATTAAATAACTTGTTCAAGGCCGCATAGCTAGTTAACAGCTGAACTAGGCTTAAAACCAACGTCTGAAGGCTCCTATTCCAGTGGCAGCTGCTGTGTGCTTCTTCTGTTTTCCATCAGTTTGGAAGGGAGCATAAAGTCTACAGCCACATGGGTGGGGTCAGCAGAAAGATTGACCACCAAGCCTGAGGCAGGTGAGGCTGATCTCCTGGGCACAGCCTCTCTGCACAGGAGTTCACAGAAGTGATATGATCCAAAGTTGCTGAGGGAAAAGCCCTTATTTGTGGAATTAACGGCAGGTCTCTCTTGAGGTCAGAATGAATGTTATTGACATTATTGTTTGTATTGTGGTAAGGTATACATAATGGAAAATGTACCATTTTGGCTGGACATGGTGGCTCACGCCTGTAATCCCAGCACTTTGGGAGGCCAAGGTGGGCGCATCACCTGAGGTCAGGAGTTCGAGACCAGCCTGCCAACATGGTGAAACCTCATCTCTACTAAAAATACAAAAATTAGCCGGGCGTGGTGGTGGGTGCCTGTAGTCCCAGCTACTTGGGAGACTGAGGCAGGAGAATCACCTGAACCCAGGAAGCAGAAG.

### Lentiviral transduction and qPCR

*ZDHHC17*-*LNCKB*.*11978* was subcloned into pCDH-CMV-Puro lentiviral vector and then co-transfected with psPAX2 and pMD2.G plasmids into HEK293T cells to generate lentiviral particles respectively. pCDH-CMV-Puro lentiviral vector was also transfected as the control. After 48 hours, the lentivirus was harvested and transduced into A549 cells with 10 μg/mL polybrene. Puromycin (1 μg/mL) was added into cells for positive selection at 48 hours post transduction for 7 days to establish stable A549 cell lines with *ZDHHC17*-*LNCKB*.*11978* fusion.

Total RNA from cells was isolated using Direct-zol RNA MiniPrep Kit (Zymo Research) according to the manufacturer’s instructions. cDNA was synthesized using SuperScript VILO cDNA synthesis kit (Life Technologies). qPCR was performed using SYBR green qPCR Master Mix (Sigma) on an Applied Biosystems QuantStudio 3 Real-Time PCR System. Primer sequences used were as follows:

*GAPDH* forward: 5’ -GTCTCCTCTGACTTCAACAGCG-3’

*GAPDH* reverse: 5’ -ACCACCCTGTTGCTGTAGCCAA-3’

*ACTIN* forward: 5’ -CACCATTGGCAATGAGCGGTTC-3’

*ACTIN* reverse: 5’ -AGGTCTTTGCGGATGTCCACGT-3’

*ZDHHC17-Inckb*.*11978* primer 1 forward: 5’ -GAGTACGATACCGAAGCGGG-3’

*ZDHHC17-Inckb*.*11978* primer 1 reverse: 5’ -ACTGAGGTGAGGAGTGGGTT-3’

*ZDHHC17-Inckb*.*11978* primer 2 forward: 5’ -CGGCCCGGATGAGTACGATA-3’

*ZDHHC17-Inckb*.*11978* primer 2 reverse: 5’ -TAACGTTCACAGCACTCGGG-3’

### Xenograft models

NOD.CB17-Prkdc^scid^/J (NOD-SCID) mice were purchased from The Jackson Laboratory. All animal experiments complied with the standards approved by University of Chicago. For tumor transplantation, 5×10^5^ A549 cells with pCDH control and *ZDHHC17*-*LNCKB*.*11978* fusions were resuspended in PBS and mixed with Matrigel (R&D Cultrex Type 3, Pathclear) at 1:1 ratio, followed by subcutaneously injection into NOD-SCID mice. Tumor volume was assessed by calipers every week. At 7 weeks post tumor grafting, animals were euthanized, and the engrafted tumors were weighed and photographed.

## Declarations

### Ethics approval and consent to participate

All animal experiments were approved by the University of Chicago IACUC and were conducted under IACUC protocol #72637. This study was carried out in strict compliance with the PHS Policy on Humane Care and Use of Laboratory Animals.

## Availability of data and materials

RNA-seq data for 9,565 tumor and 715 normal samples from The Cancer Genome Atlas (TCGA) (**Supplementary Tables S1**) were downloaded from Genomic Data Commons (https://portal.gdc.cancer.gov/). RNA-seq data for MCF7, HCT116, and K562 cell lines were downloaded from the National Center for Biotechnology Information (NCBI) Sequence Read Archive (SRA) with accession SRX5414642 (MCF7, CCLE), SRX159831 (MCF7, ENCODE), SRX6378523 (MCF7 Weber et al.), SRX6378524 (MCF7 Weber et al.), SRX5414471 (HCT116, CCLE) and SRX159835 (HCT116, ENCODE), SRX5414683 (K562, CCLE), SRX1603406 (K562, ENCODE) and SRX1603407 (K562, ENCODE). RNA-seq data for two normal adipose tissue samples from Genotype-Tissue Expression (GTEx) were downloaded from NCBI SRA with accession SRX636240 and SRX640265.

Somatic SVs in TCGA samples were obtained from a recent Pan-cancer Analysis of Whole Genomes (PCAWG) study^25^. Somatic SVs in MCF7 were downloaded from the Dependency Map (DepMap) portal (https://depmap.org/portal/). Fusions in TCGA samples identified by Arriba, DEEPEST, and STAR-Fusion were downloaded from the related publications^3,12,16^. Fusions in MCF7 identified by FusionCatcher (v1.0), InFusion (v0.8), MapSplic2 (v2.2.1), SOAPfuse (v1.2.7), and STAR-Fusion (v1.5.0) were downloaded from the previous study^24^. Fusions in MCF7 identified by EasyFuse (v1.3.0) were provided by Dr. Ugur Sahin. The subtypes of sarcomas were obtained from a previous study^31^.

All coordinates were based on hg38 reference genome. GENCODE v29 was used for gene annotation. NOCODE v6 and lncRNAKB v7 were used to annotate non-coding genes that are not annotated by GENOCDE.

## Availability of software

The SFyNCS package is available at https://github.com/yanglab-computationalgenomics/SFyNCS.

## Author contributions

Software, X.Z., and L.Y.; analysis, X.Z. and L.Y.; PCR and Sanger sequencing: X.Z., A.Y., and A.LH.; animal experiment, J.L., A.Y., Y.M. and L.Y.; conceptulation, L.Y.; writing, L.Y.; supervision, Y.M. and L.Y. All authors have read and approved the final manuscript.

## References

1. Mitelman, F., Johansson, B. & Mertens, F. The impact of translocations and gene fusions on cancer causation. Nat. Rev. Cancer 7, 233–245 (2007).

2. Mertens, F., Johansson, B., Fioretos, T. & Mitelman, F. The emerging complexity of gene fusions in cancer. Nat. Rev. Cancer 15, 371–381 (2015).

3. Gao, Q. et al. Driver Fusions and Their Implications in the Development and Treatment of Human Cancers. Cell Rep. 23, 227–238.e3 (2018).

4. Savage, D. G. & Antman, K. H. Imatinib Mesylate — A New Oral Targeted Therapy. N. Engl. J. Med. 346, 683–693 (2002).

5. Schram, A. M., Chang, M. T., Jonsson, P. & Drilon, A. Fusions in solid tumours: Diagnostic strategies, targeted therapy, and acquired resistance. Nature Reviews Clinical Oncology vol. 14 735–748 (2017).

6. Jang, Y. E. et al. ChimerDB 4.0: An updated and expanded database of fusion genes. Nucleic Acids Res. 48, D817–D824 (2020).

7. Tomlins, S. A. et al. Distinct classes of chromosomal rearrangements create oncogenic ETS gene fusions in prostate cancer. Nature 448, 595–599 (2007).

8. Nakamura, Y. et al. The GAS5 (growth arrest-specific transcript 5) gene fuses to BCL6 as a result of t(1;3)(q25;q27) in a patient with B-cell lymphoma. Cancer Genet. Cytogenet. 182, 144–149 (2008).

9. Ren, S. et al. RNA-seq analysis of prostate cancer in the Chinese population identifies recurrent gene fusions, cancer-associated long noncoding RNAs and aberrant alternative splicings. Cell Res. 22, 806–821 (2012).

10. Spans, L. et al. Recurrent MALAT1–GLI1 oncogenic fusion and GLI1 up-regulation define a subset of plexiform fibromyxoma. J. Pathol. 239, 335–343 (2016).

11. Kleinman, C. L. et al. Fusion of TTYH1 with the C19MC microRNA cluster drives expression of a brain-specific DNMT3B isoform in the embryonal brain tumor ETMR. Nat. Genet. 46, 39–44 (2014).

12. Guo, M. et al. The landscape of long noncoding RNA-involved and tumor-specific fusions across various cancers. Nucleic Acids Res. 48, 12618–12631 (2020).

13. Hayes, J., Peruzzi, P. P. & Lawler, S. MicroRNAs in cancer: biomarkers, functions and therapy. Trends Mol. Med. 20, 460–469 (2014).

14. Li, Z. & Rana, T. M. Therapeutic targeting of microRNAs: current status and future challenges. Nat. Rev. Drug Discov. 13, 622–638 (2014).

15. Nussbacher, J. K., Tabet, R., Yeo, G. W. & Lagier-Tourenne, C. Disruption of RNA Metabolism in Neurological Diseases and Emerging Therapeutic Interventions. Neuron vol. 102 294–320 (2019).

16. Dehghannasiri, R. et al. Improved detection of gene fusions by applying statistical methods reveals oncogenic RNA cancer drivers. Proc. Natl. Acad. Sci. U. S. A. 116, 15524–15533 (2019).

17. Uhrig, S. et al. Accurate and efficient detection of gene fusions from RNA sequencing data. Genome Res. 31, 448–460 (2021).

18. Nicorici, D. et al. FusionCatcher – a tool for finding somatic fusion genes in paired-end RNA-sequencing data. bioRxiv 011650 (2014) doi:10.1101/011650.

19. Okonechnikov, K. et al. InFusion: Advancing Discovery of Fusion Genes and Chimeric Transcripts from Deep RNA-Sequencing Data. PLoS One 11, (2016).

20. McPherson, A. et al. deFuse: An Algorithm for Gene Fusion Discovery in Tumor RNA-Seq Data. PLoS Comput. Biol. 7, 1001138 (2011).

21. Ma, C., Shao, M. & Kingsford, C. SQUID: Transcriptomic structural variation detection from RNA-seq. Genome Biol. 19, 1–16 (2018).

22. Wang, K. et al. MapSplice: accurate mapping of RNA-seq reads for splice junction discovery. Nucleic Acids Res. 38, e178–e178 (2010).

23. Jia, W. et al. SOAPfuse: An algorithm for identifying fusion transcripts from paired-end RNA-Seq data. Genome Biol. 14, 1–15 (2013).

24. Weber, D. et al. Accurate detection of tumor-specific gene fusions reveals strongly immunogenic personal neo-antigens. Nat. Biotechnol. 2022 408 40, 1276–1284 (2022).

25. Li, Y. et al. Patterns of somatic structural variation in human cancer genomes. Nature 578, 112–121 (2020).

26. Yang, L. et al. Analyzing Somatic Genome Rearrangements in Human Cancers by Using Whole-Exome Sequencing. Am. J. Hum. Genet. 98, 843–856 (2016).

27. Campbell, P. J. et al. Pan-cancer analysis of whole genomes. Nature 578, 82–93 (2020).

28. Dobin, A. et al. STAR: ultrafast universal RNA-seq aligner. Bioinformatics 29, 15–21 (2013).

29. Kim, D. et al. TopHat2: Accurate alignment of transcriptomes in the presence of insertions, deletions and gene fusions. Genome Biol. 14, 1–13 (2013).

30. Kent, W. J. BLAT—The BLAST-Like Alignment Tool. Genome Res. 12, 656 (2002).

31. Abeshouse, A. et al. Comprehensive and Integrated Genomic Characterization of Adult Soft Tissue Sarcomas. Cell 171, 950–965.e28 (2017).

