## Supplementary figures for "SFyNCS detects oncogenic fusions involving non-coding sequences in cancer"

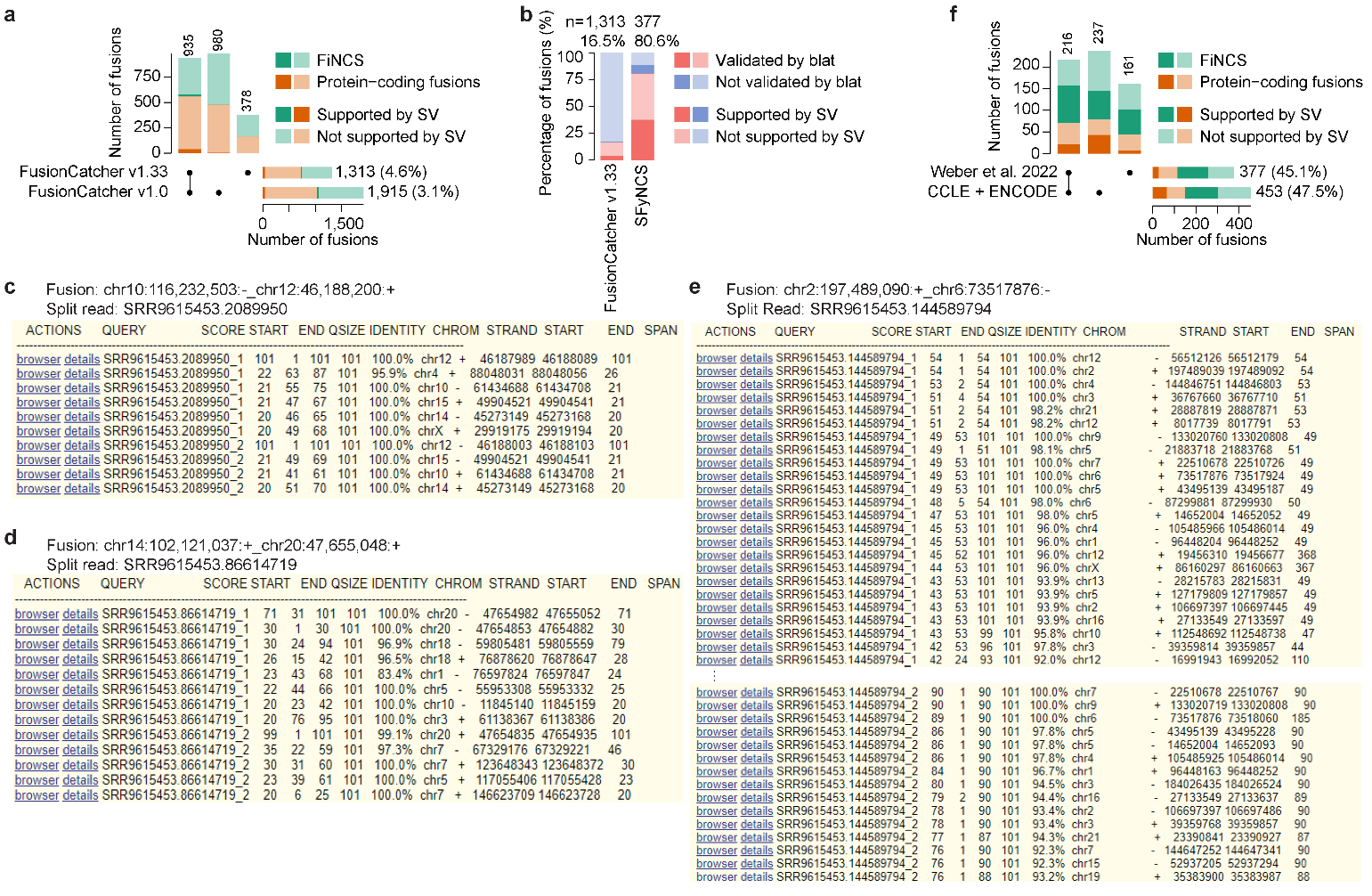

**Figure S1. Benchmarking fusion detection in MCF7 cell line. a**, Comparison between FusionCatcher v1.0 and v1.33. The stacked bars on the top show the number of fusions identified by one or both versions. The stacked bars on the bottom right are the total fusions detected by the two versions. The black dots under the stacked bars indicate versions used. The numbers on the top and on the right side of the bars are numbers of fusions. The percentages in the parenthesis indicate percentages of fusions supported by somatic SVs. **b**, Barplot showing re-alignments of split reads by BLAT for fusions detected by FusionCatcher v1.33 and SFyNCS. **c**, **d** and **e**, Examples of BLAT alignments that do not support the predicted fusions. Coordinates of predicted fusions and read names of split reads are listed on the top. BLAT screenshots are provided for split reads aligned to the hg38 reference genome. **c**, The split read can be aligned entirely to one location of the genome. **d**, The split read only supports one fusion breakpoint (chr20:47,655,048:+) but not the other (chr14:102,121,037:+). **e**, The split read can be aligned to many genomic regions. There are 105 alignments provided by BLAT and only 39 are shown. **f**, Comparison between fusions detected by SFyNCS from RNAseq data produced by Weber et al. 2022 as well as CCLE and ENCODE.

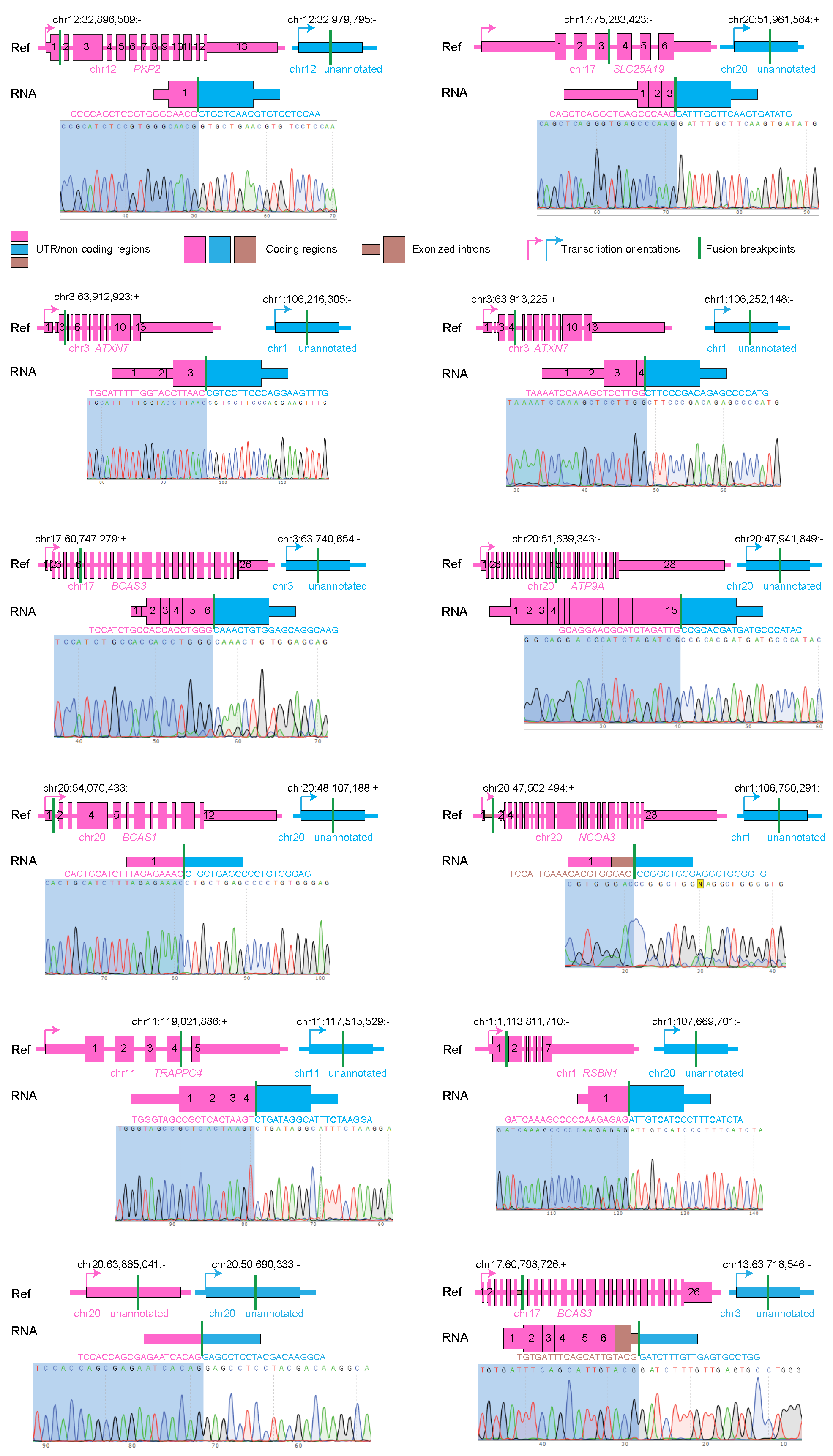

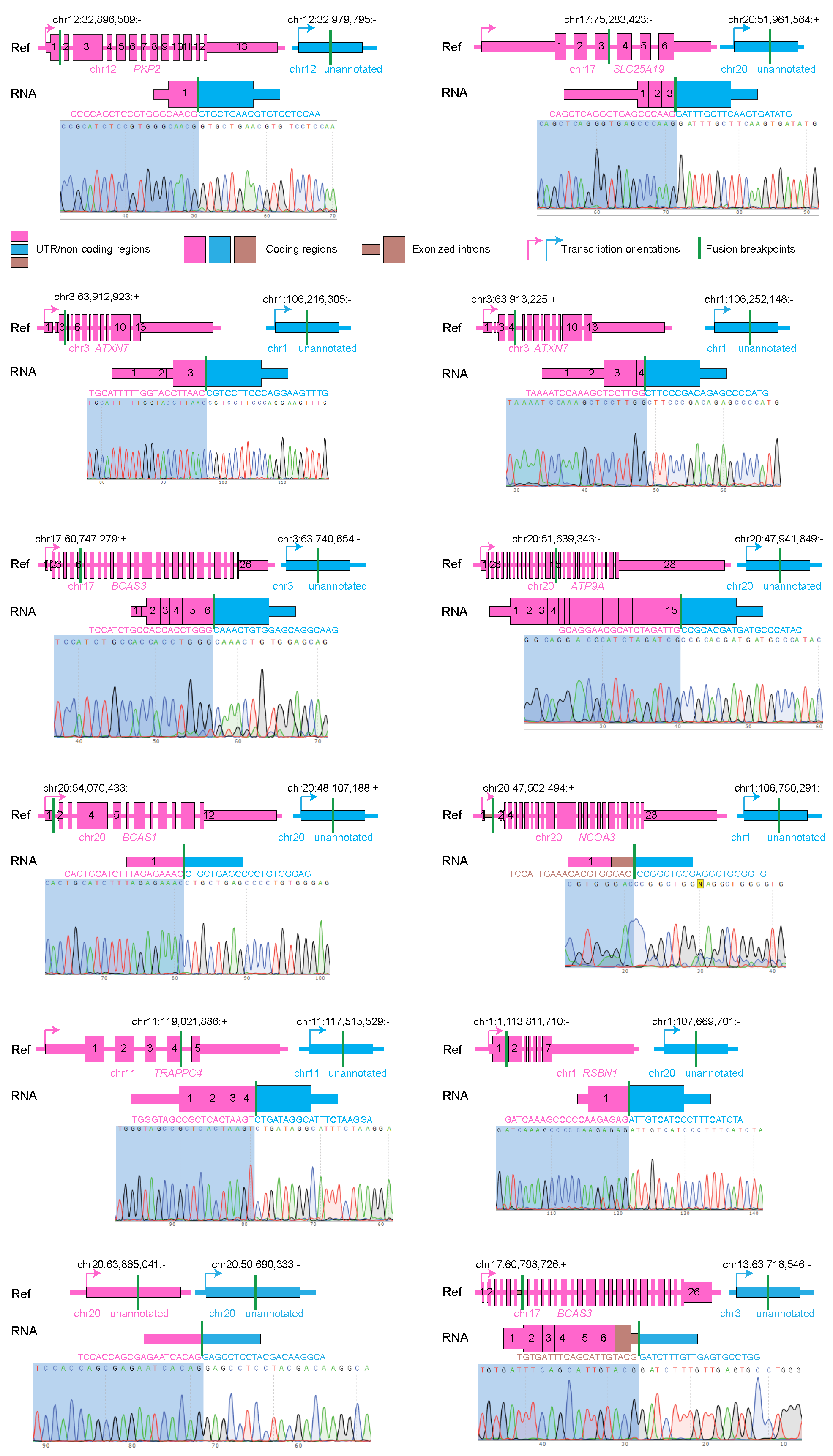

**Figure S2. Sanger sequencing validation for FiNCS in cell line MCF7 only detected by SFyNCS, but not by STAR-Fusion, MapSplice2, InFusion, SOAPfuse, EasyFuse or FusionCatcher.** Gene structures are shown in the reference tracks and fusion structures are shown in the fusion tracks. Sequences at the fusion breakpoints are shown above the Sanger sequencing chromatograms.

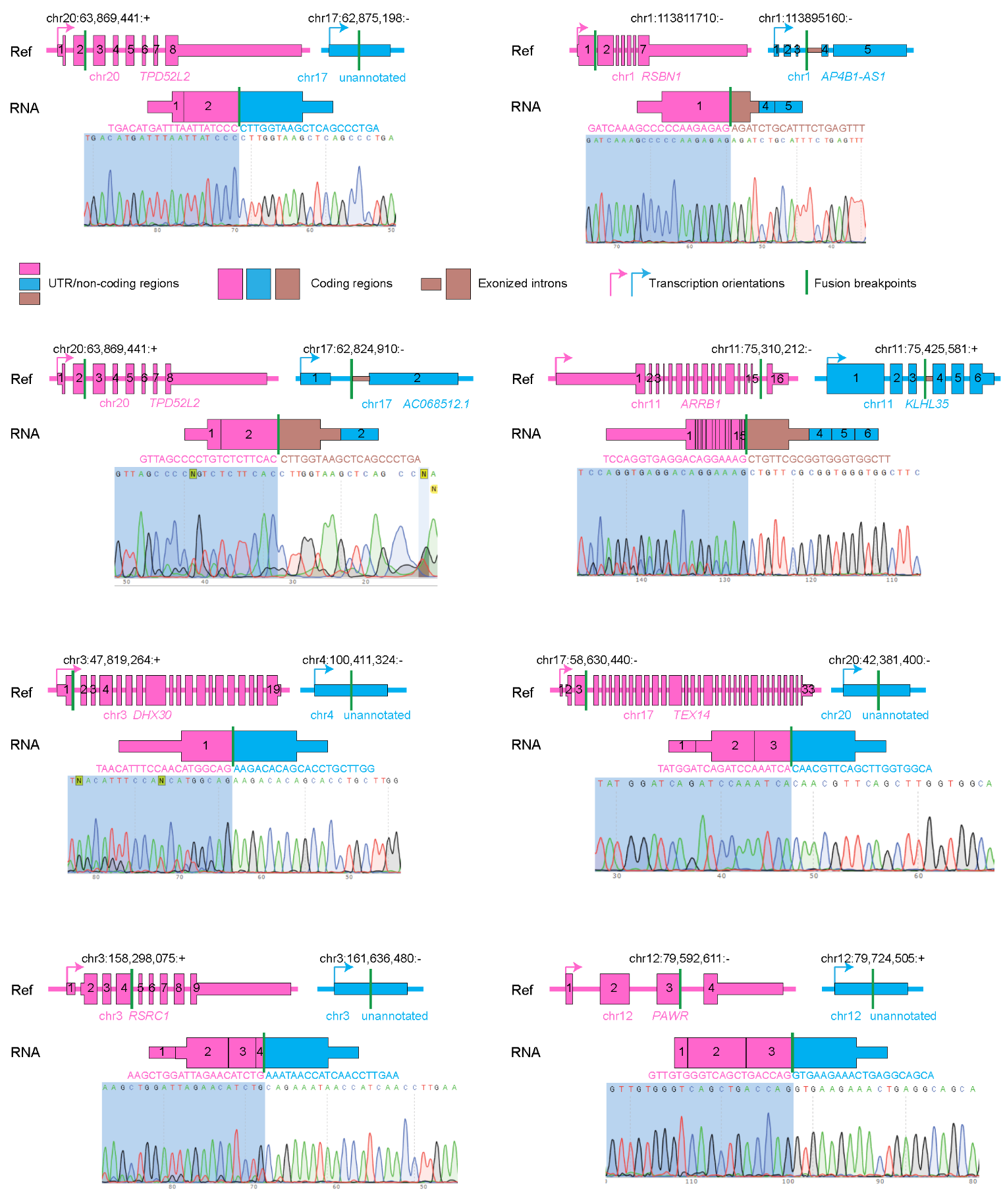

**Figure S3. Sanger sequencing validation for FiNCS in cell line MCF7 only detected from RNAseq data from CCLE and ENCODE but not in Weber et al. 2022.** Gene structures are shown in the reference tracks and fusion structures are shown in the fusion tracks. Sequences at the fusion breakpoints are shown above the Sanger sequencing chromatograms.

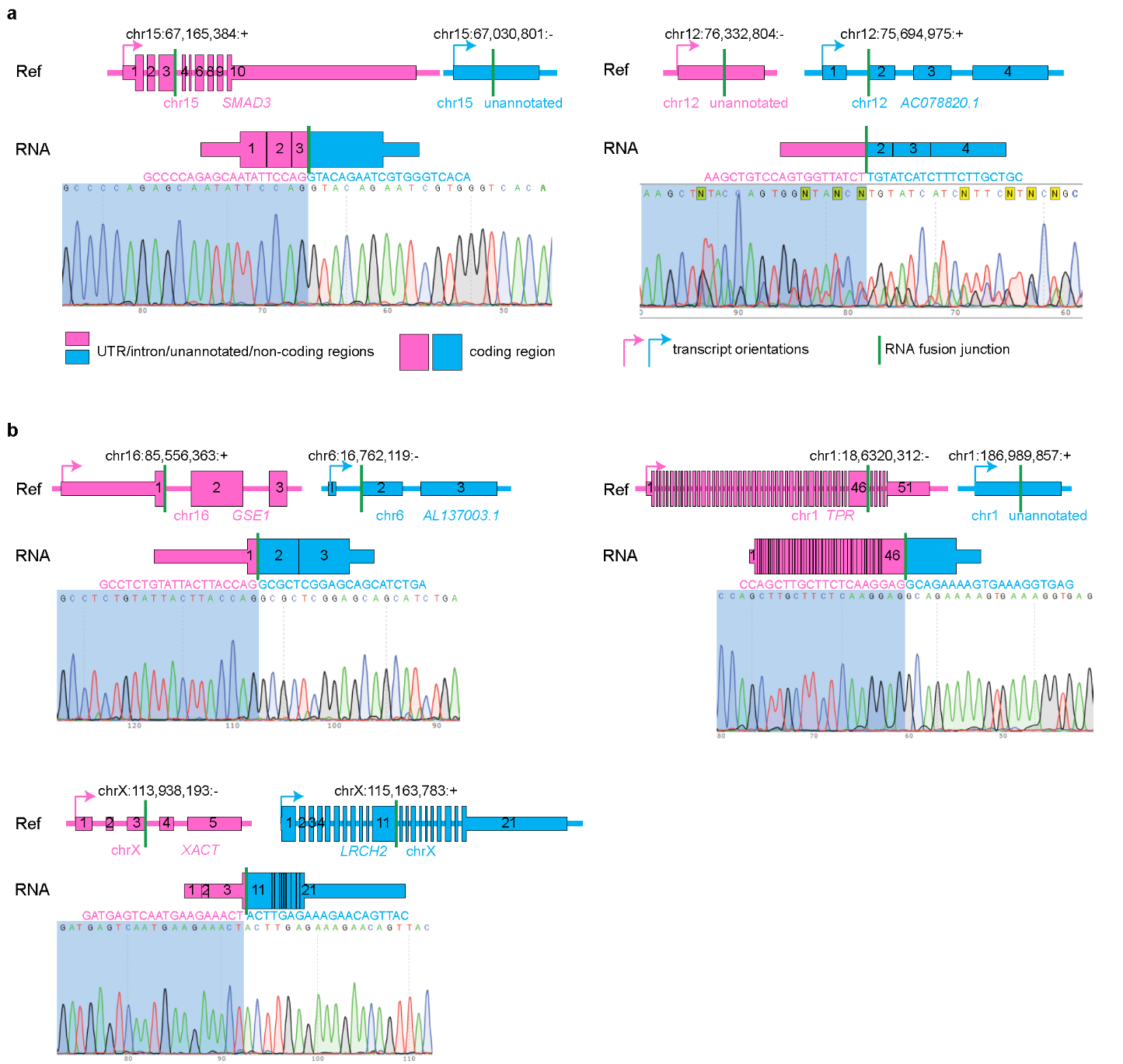

**Figure S4. Sanger sequencing validation for FiNCS in cell lines HCT116 and K562.** **a**, FiNCS in HCT116. **b**, FiNCS in K562. Gene structures are shown in the reference tracks and fusion structures are shown in the fusion tracks. Sequences at the fusion breakpoints are shown above the Sanger sequencing chromatograms.

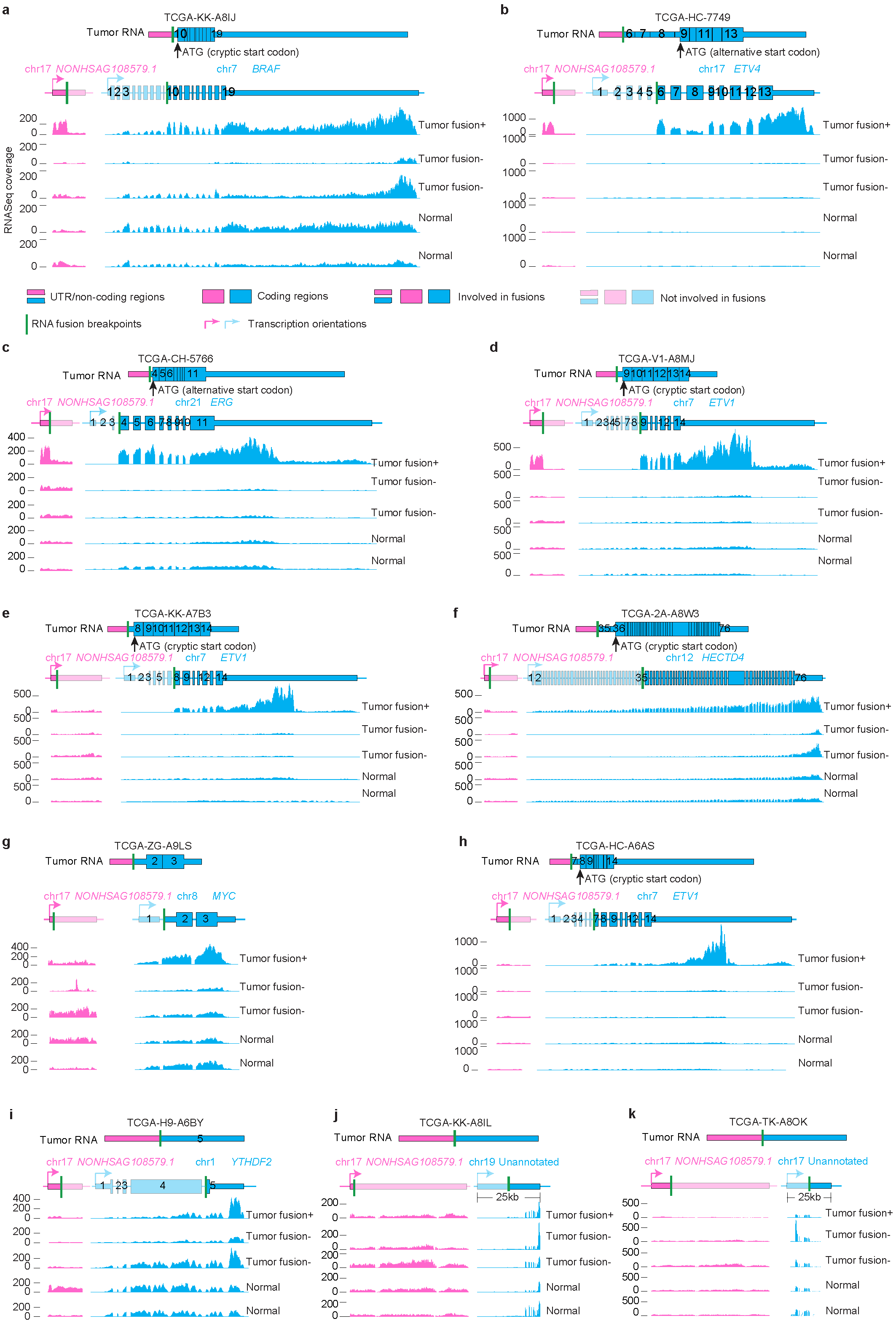

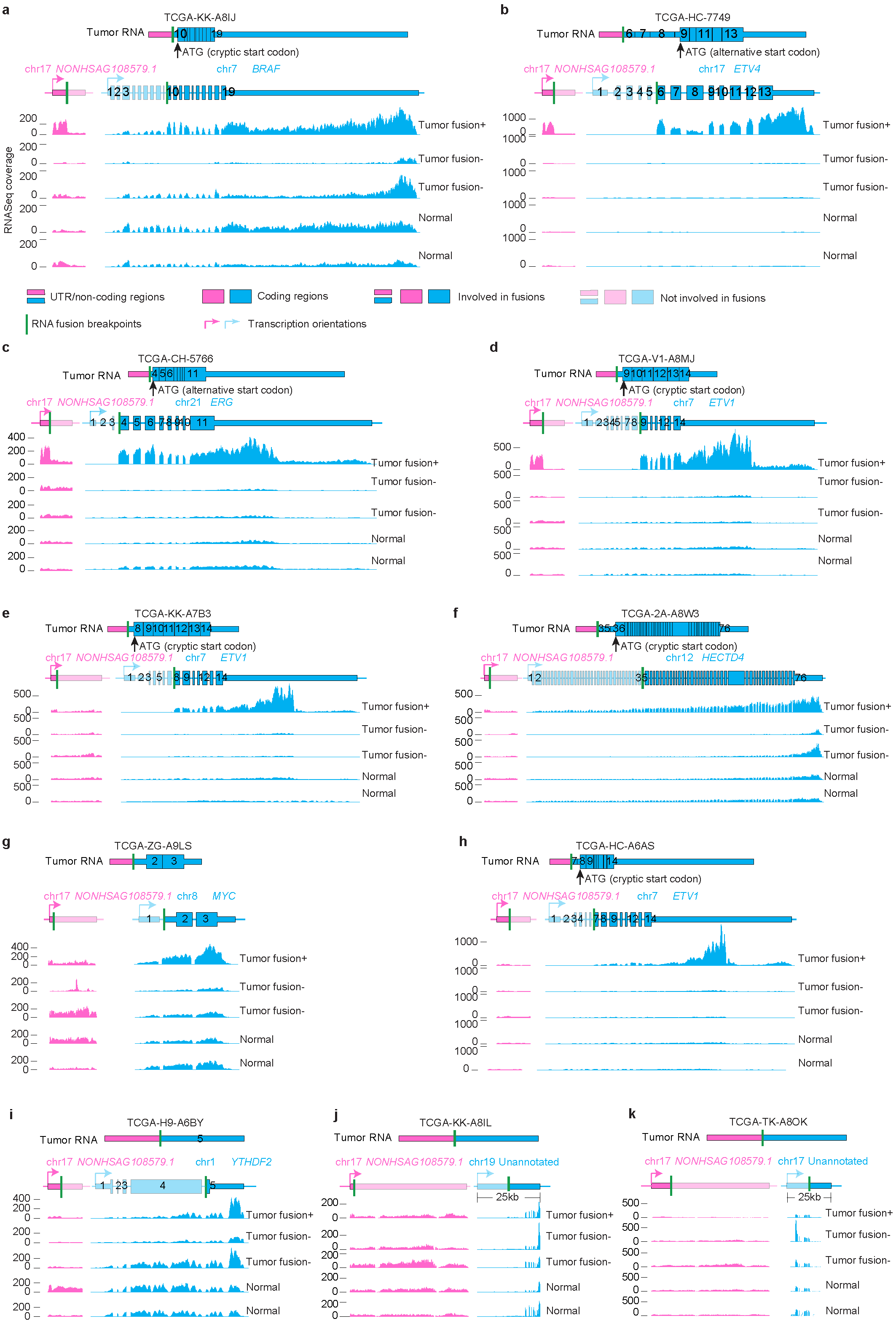

**Figure S5. Additional *NONHSAG108579.1* fusions in prostate cancers and their expression.** Fusion structures are given on the top. Five tracks of RNAseq coverage are shown for five samples at the bottom and the reference gene structures are given above the five tracks. Exons and introns are re-scaled to better illustrate fusion structures. In **a**, the tumor samples without fusions (fusion-) are TCGA-HI-7169-01A-11R-2118-07 and TCGA-G9-6365-01A-11R-1789-07, the normal samples are TCGA-EJ-7123-11A-01R-1965-07 and TCGA-EJ-7125-11A-01R-1965-07. In **b**, the fusion- samples are TCGA-HI-7169-01A-11R-2118-07 and TCGA-G9-6365-01A-11R-1789-07, the normal samples are TCGA-EJ-7123-11A-01R-1965-07 and TCGA-EJ-7125-11A-01R-1965-07. In **c**, the fusion- samples are TCGA-FC-A5OB-01A-11R-A29R-07 and TCGA-V1-A9OL-01A-11R-A41O-07, the normal samples are TCGA-EJ-A8FO-11A-11R-A36G-07 and TCGA-EJ-7331-11A-01R-2118-07. In **d**, the fusion- samples are TCGA-G9-6365-01A-11R-1789-07 and TCGA-V1-A9OL-01A-11R-A41O-07, the normal samples are TCGA-EJ-A8FO-11A-11R-A36G-07 and TCGA-EJ-7331-11A-01R-2118-07. In **e**, the fusion- samples are TCGA-G9-6365-01A-11R-1789-07 and TCGA-V1-A9OL-01A-11R-A41O-07, the normal samples are TCGA-EJ-A8FO-11A-11R-A36G-07 and TCGA-EJ-7331-11A-01R-2118-07. In **f**, the fusion- samples are TCGA-HI-7169-01A-11R-2118-07 and TCGA-G9-6365-01A-11R-1789-07, the normal samples are TCGA-EJ-A8FO-11A-11R-A36G-07 and TCGA-EJ-7331-11A-01R-2118-07. In **g**, the fusion- samples are TCGA-HI-7169-01A-11R-2118-07 and TCGA-EJ-A7NJ-01A-22R-A352-07, the normal samples are TCGA-EJ-7327-11A-01R-2118-07 and TCGA-HC-7742-11A-01R-2118-07. In **h**, the fusion- samples are TCGA-G9-6365-01A-11R-1789-07 and TCGA-V1-A9OL-01A-11R-A41O-07, the normal samples are TCGA-EJ-A8FO-11A-11R-A36G-07 and TCGA-EJ-7331-11A-01R-2118-07. In **i**, the fusion- samples are TCGA-HI-7169-01A-11R-2118-07 and TCGA-G9-6365-01A-11R-1789-07, the normal samples are TCGA-EJ-7327-11A-01R-2118-07 and TCGA-EJ-7331-11A-01R-2118-07. In **j**, the fusion- samples are TCGA-G9-6365-01A-11R-1789-07 and TCGA-V1-A9OL-01A-11R-A41O-07, the normal samples are TCGA-EJ-A8FO-11A-11R-A36G-07 and TCGA-EJ-7331-11A-01R-2118-07. In **k**, the fusion- samples are TCGA-G9-6365-01A-11R-1789-07 and TCGA-V1-A9OL-01A-11R-A41O-07, the normal samples are TCGA-EJ-A8FO-11A-11R-A36G-07 and TCGA-EJ-7331-11A-01R-2118-07.

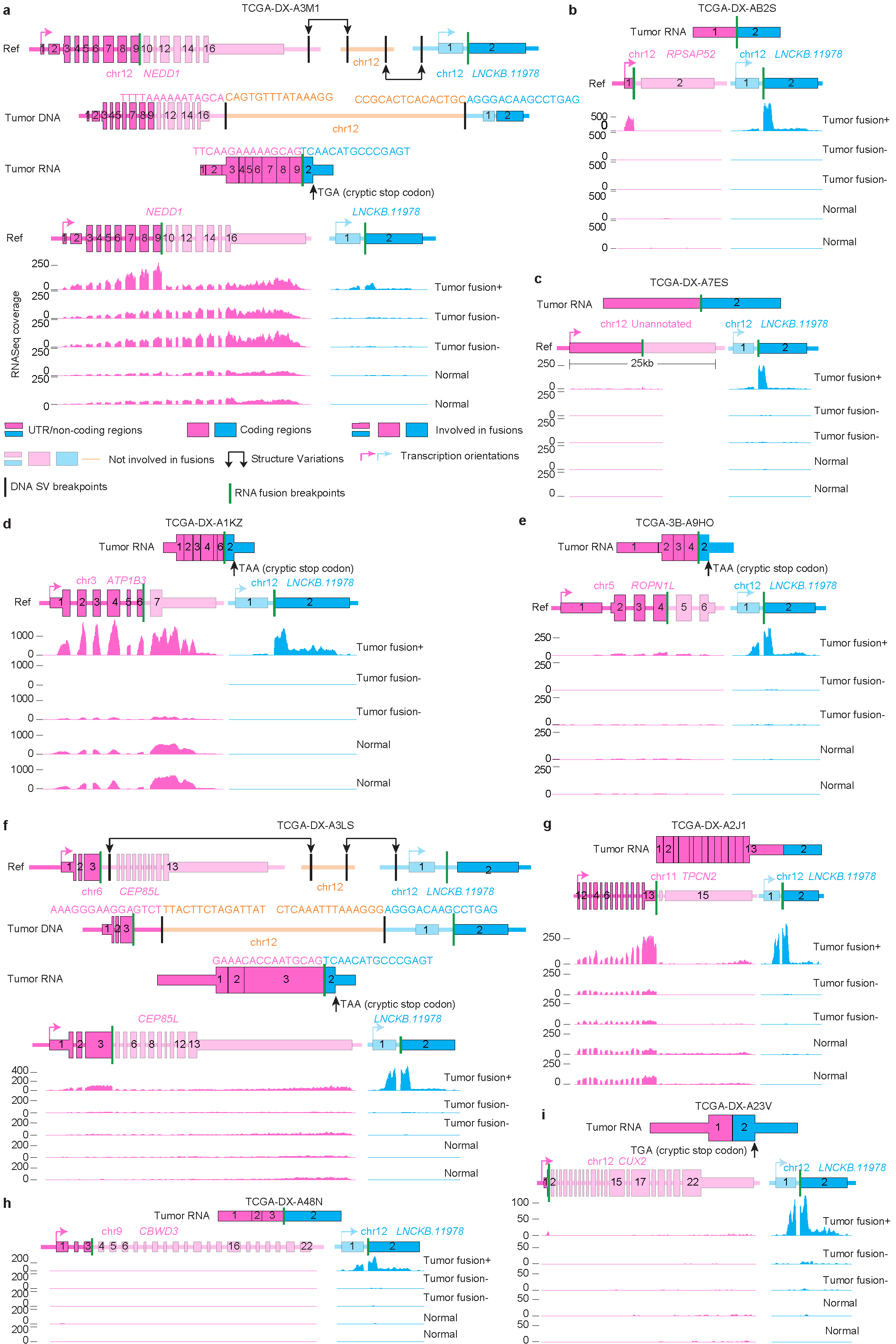

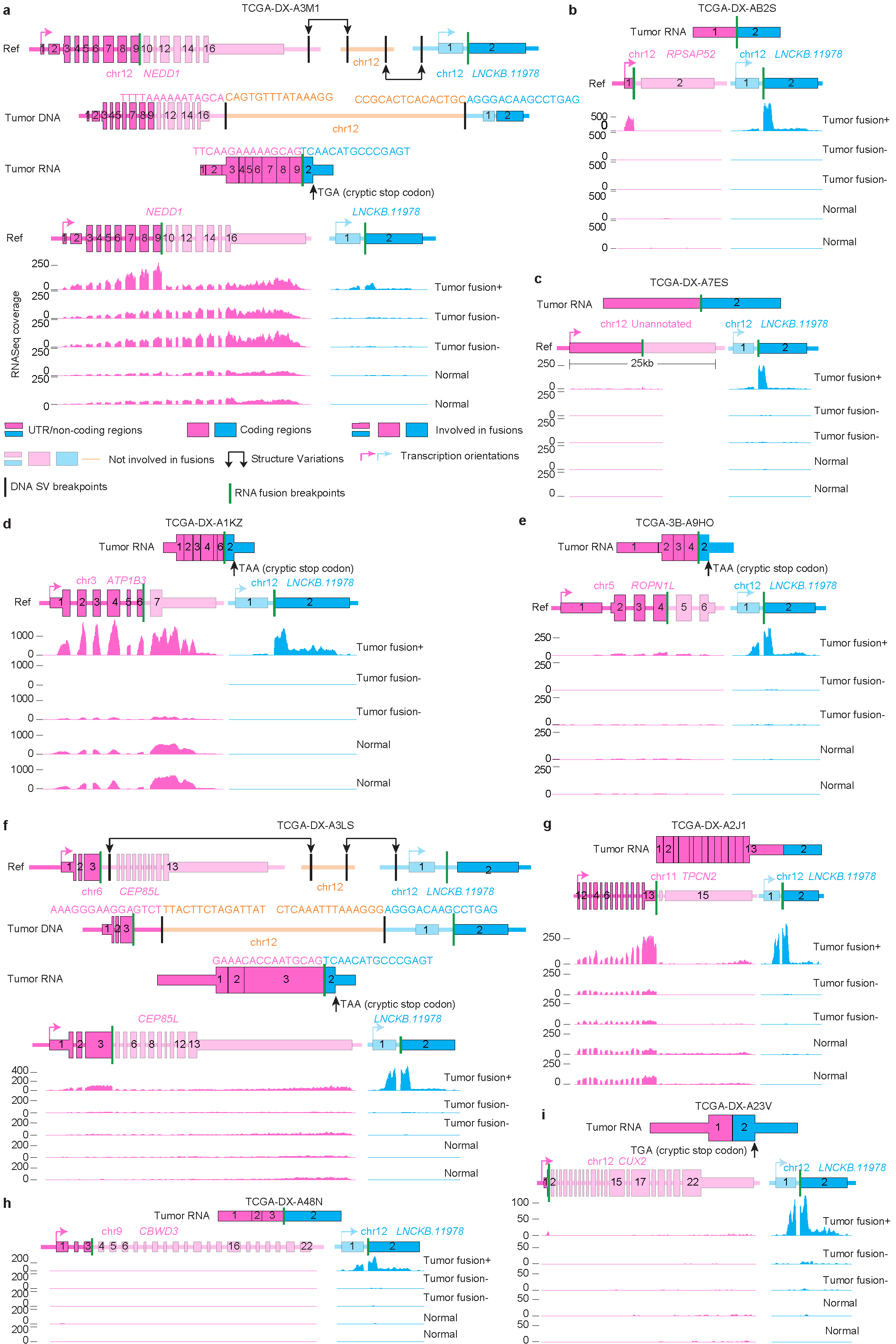

**Figure S6.** **Additional *LNCKB.11978*** **fusions in DDLPS and their expression. a** and **f**, Samples with WGS and RNAseq. Gene and fusion structures of reference genome, tumor DNA and tumor RNA are shown as the top three tracks. In all cases, five tracks of RNAseq coverage are shown for five samples at the bottom and the reference gene structures are given above the five tracks. Exons and introns are re-scaled to better illustrate fusion structures. In **a**-**i**, the tumor samples without fusions (fusion-) are TCGA-IE-A4EI-01A-11R-A24X-07 and TCGA-IW-A3M4-01A-11R-A21T-07, the normal samples are SRR1485722 and SRR1498838.

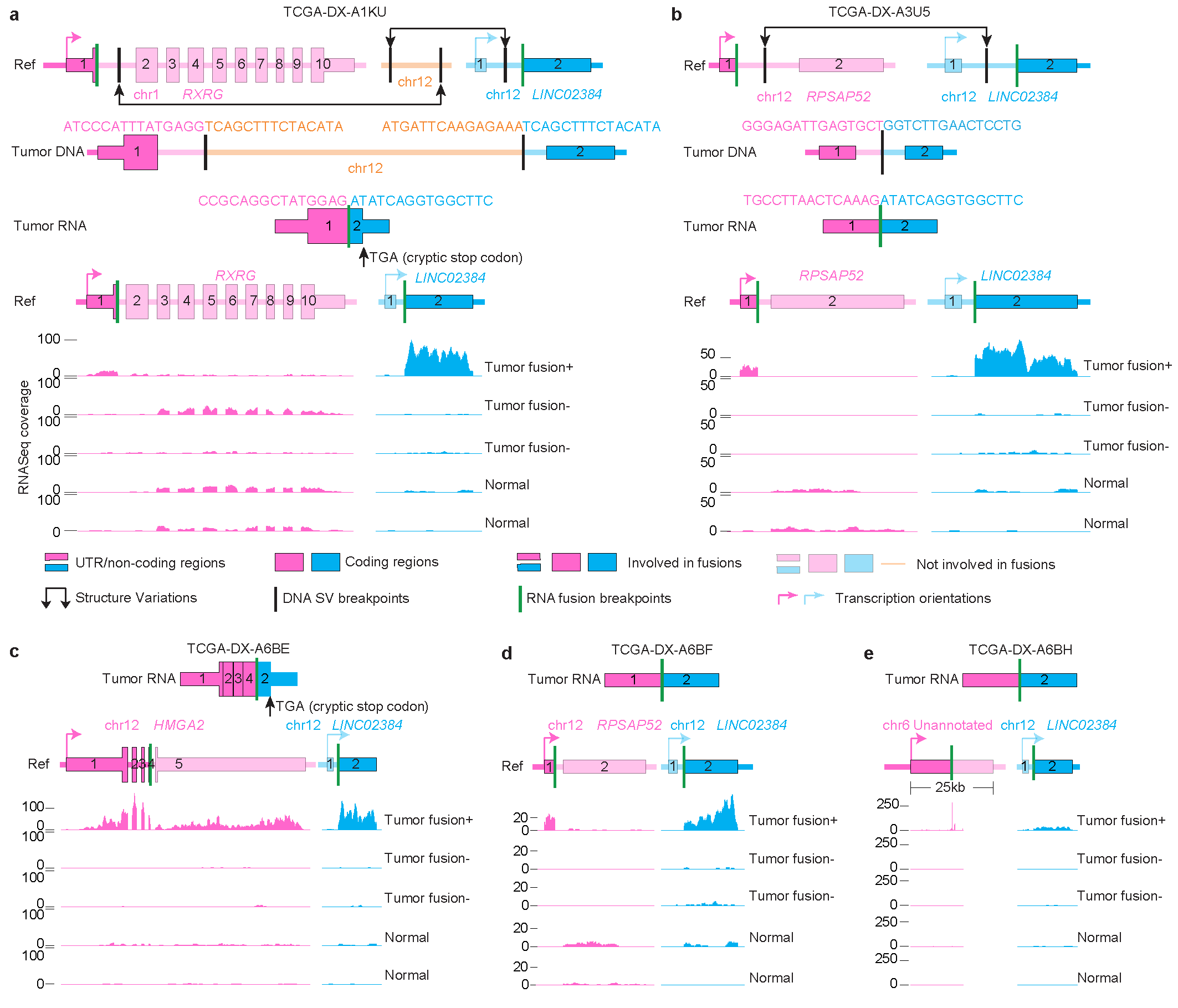

**Figure S7.** **Additional *LINC02384*** **fusions in DDLPS and their expression. a** and **b**, Samples with WGS and RNAseq. Gene and fusion structures of reference genome, tumor DNA and tumor RNA are shown as the top three tracks. In all cases, five tracks of RNAseq coverage are shown for five samples at the bottom and the reference gene structures are given above the five tracks. Exons and introns are re-scaled to better illustrate fusion structures. In **a**-**e**, the tumor samples without fusions (fusion-) are TCGA-IE-A4EI-01A-11R-A24X-07 and TCGA-IW-A3M4-01A-11R-A21T-07, the normal samples are SRR1485722 and SRR1498838.

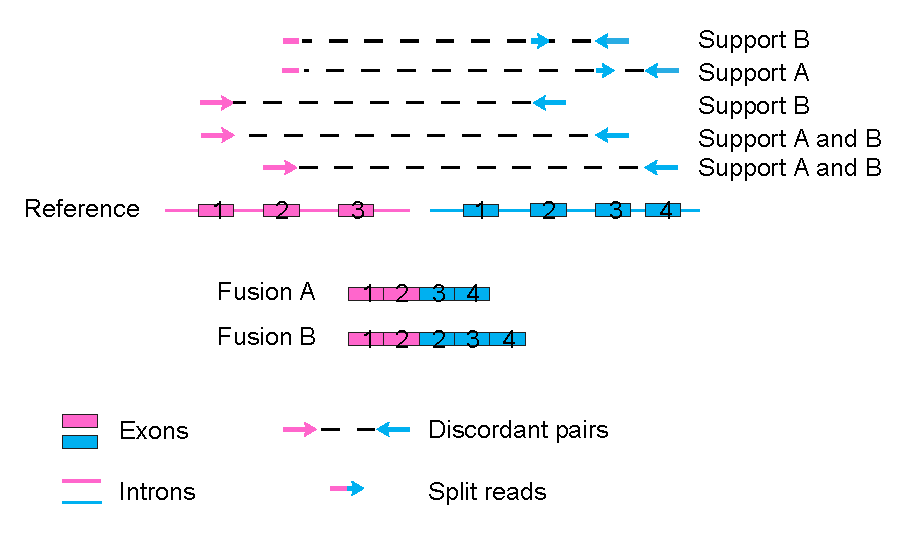

**Figure S8. Scheme of discordant read pairs and split reads supporting different isoforms.** Two isoforms are produced from the fusion locus. Some discordant pairs can support multiple isoforms.
